# Rapid Report: Retinal gene and pathway modulation by benzathine penicillin: A potential link to Extensive Macular Atrophy with Pseudodrusen (EMAP) and neurodegeneration

**DOI:** 10.1101/2025.09.15.676285

**Authors:** Rogério N. Shinsato, Roberto H. Herai

**Affiliations:** Unisalesiano, Araçatuba, São Paulo, 16016-500, Brazil; Laboratory of Bioinformatics and Neurogenetics (LaBiN/LEM), Graduate Program in Health Sciences, School of Medicine and Life Sciences, Pontifícia Universidade Católica do Paraná (PUCPR), Curitiba, Paraná, 80215-901, Brazil; Research Division, Buko Kaesemodel Institute (IBK), Curitiba, Paraná, 80240-000, Brazil; Research Division, 9p Brazil Association (A9pB), Santa Maria, Rio Grande do Sul, 97060-580, Brazil

**Keywords:** Penicillin benzathine, neurodegeneration, retina, genetic modulation

## Abstract

Extensive Macular Atrophy with Pseudodrusen (EMAP) is a rare and rapidly progressive retinal disease marked by central vision loss, chorioretinal atrophy, and pseudodrusen, primarily affecting middle-aged individuals. Recent studies have identified a statistically significant association between long-term use of benzathine penicillin (BP) and the onset of EMAP. However, the molecular mechanisms underlying this relationship remain unclear. In this study, we performed an integrative in silico analysis to explore the genes modulated by BP that are expressed in ocular tissues and involved in retinal metabolic pathways. Using public databases (PubChem, GTEx, HPA, EYEDB), we identified 52 BP-responsive genes, with six (*CD4, CRP, IL6, IL1R1, TAC1, TNF*) directly modulated by BP and expressed in the retina. These genes are implicated in inflammation, immune response, and retinal homeostasis. Expression data show strong presence in eye and brain tissues, and several are associated with age-related macular degeneration (AMD). Network analysis revealed gene–gene and protein–protein interactions, highlighting a shared pathophysiological axis. Notably, IL6 and TNF are involved in oxidative stress, endoplasmic reticulum stress, and chronic inflammation—hallmarks of both EMAP and atrophic AMD. Our findings suggest that BP may induce EMAP by dysregulating retinal genes involved in neurodegeneration, providing new insights into drug safety and retinal disease mechanisms.

## Main text

Extensive Macular Atrophy with Pseudodrusen (EMAP) is a rapidly progressive form of macular atrophy with pseudodrusen characteristics that appear in middle-aged individuals (Hamel, Meunier et al. 2009).

EMAP also includes the presence of pavinng stone-like alterations in the retina periphery and chorioretinal atrophy that extends to the arcades (Carlà, Giannuzzi et al. 2024). Affecting more women, initial symptoms of EMAP include a complaint of night blindness and rapidly progressing to loss of central vision (Antropoli, Bianco et al. 2024). Clinically, patients with EMAP disease have elevated plasma levels of C3 and decreased levels of CH50 proteins (Douillard, Picot et al. 2016), and its genetic findings are still limited to weak associations involving mutations in the genes *C1QTNF5* and *TIMP3* (Antropoli, Bianco et al. 2024).

Recently, rapidly emerged evidence indicated a statistically significant correlation between prolonged use of benzathine penicillin (BP) as the cause of EMAP. This drug-disease association was identified by a cohort study (formed by 77 patients with EMAP) that investigated the use of BP to treat rheumatic fever (Moreira-Neto et al. 2024). Over decades, BP has been widely used as an intramuscular drug treatment to reduce the risk of rheumatic fever recurrences (Bray, Thompson et al. 2024). Although the recent findings indicate a significant connection between BP as the factual cause of EMAP, it is still unclear how the drug can cause molecular modifications that can drive pathophysiological alterations in retina. Thus, understanding the molecular cascade associated with the use of BP can provide valuable data on drug safety, its harmful effects and some mechanistic understanding of the EMAP disease.

Here, we performed a critical analysis to show, for the first time, how the drug BP can cause molecular modifications in retina that can lead to EMAP. We identified all genes that can be directly or indirectly modulated by BP. Next, we isolated only those genes related to eye and found 6 genes that can be directly associated with EMAP. We verified the expression of these eye-related genes in distinct parts of the body, including the brain and the eye. Finally, we also performed an integrative analysis to investigate the role of the EMAP-related genes in the retina by discussing their association with different pathways and how misregulation of the genes can lead to EMAP.

To identify all the human genes that can be modulated (direct or indirectly) by BP, we used the PubChem databank (Kim, Chen et al. 2025) and found 52 associated genes (Table1). The list of genes modulated by BP were found by independent studies that found several biological processes affected by this drug. The genes BMP3 and BMPR1B are involved in in bone repair and regeneration (Cui et al., 2018; Giovanini et al., 2018). The genes *CNR1, ENPEP, ERVW-1, HTT, IL1R1, NPS, PAH, RPA1, TAC1, TFCP2 and TH* were linked to BP by causing neurological responses (Table 1). Moreover, 6 out of the 52 genes (CD4, CRP, IL6, TNF, IL1R1 and TAC1) are described to be directly affected by BP with variable evidence level (Table 2).

**Table 1.**
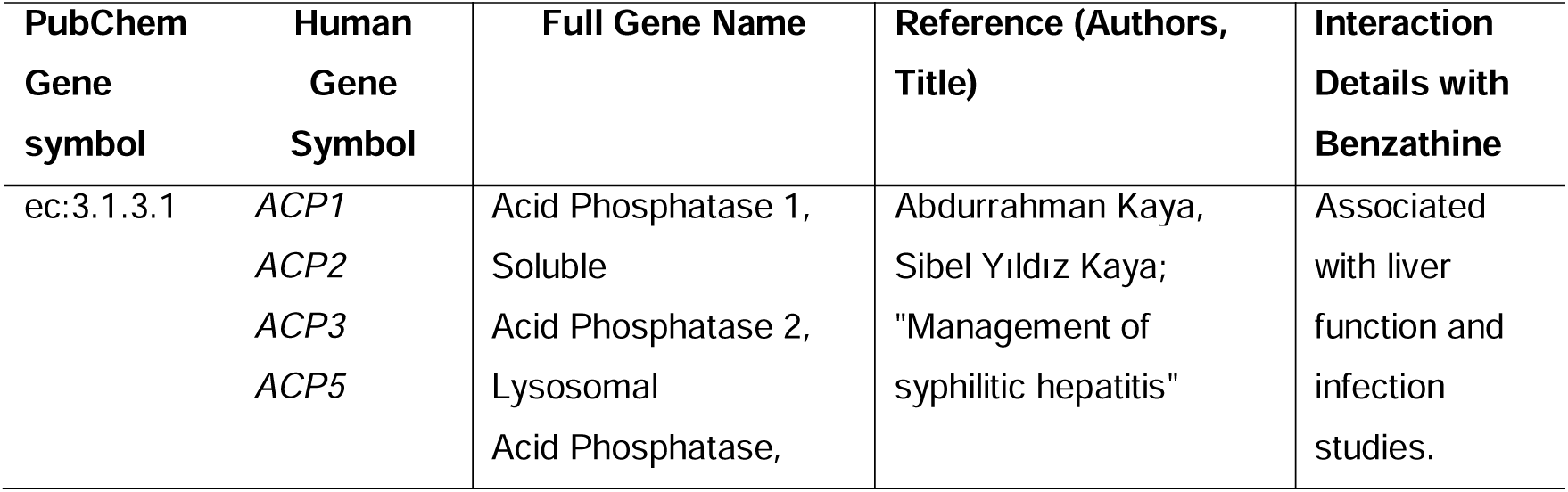
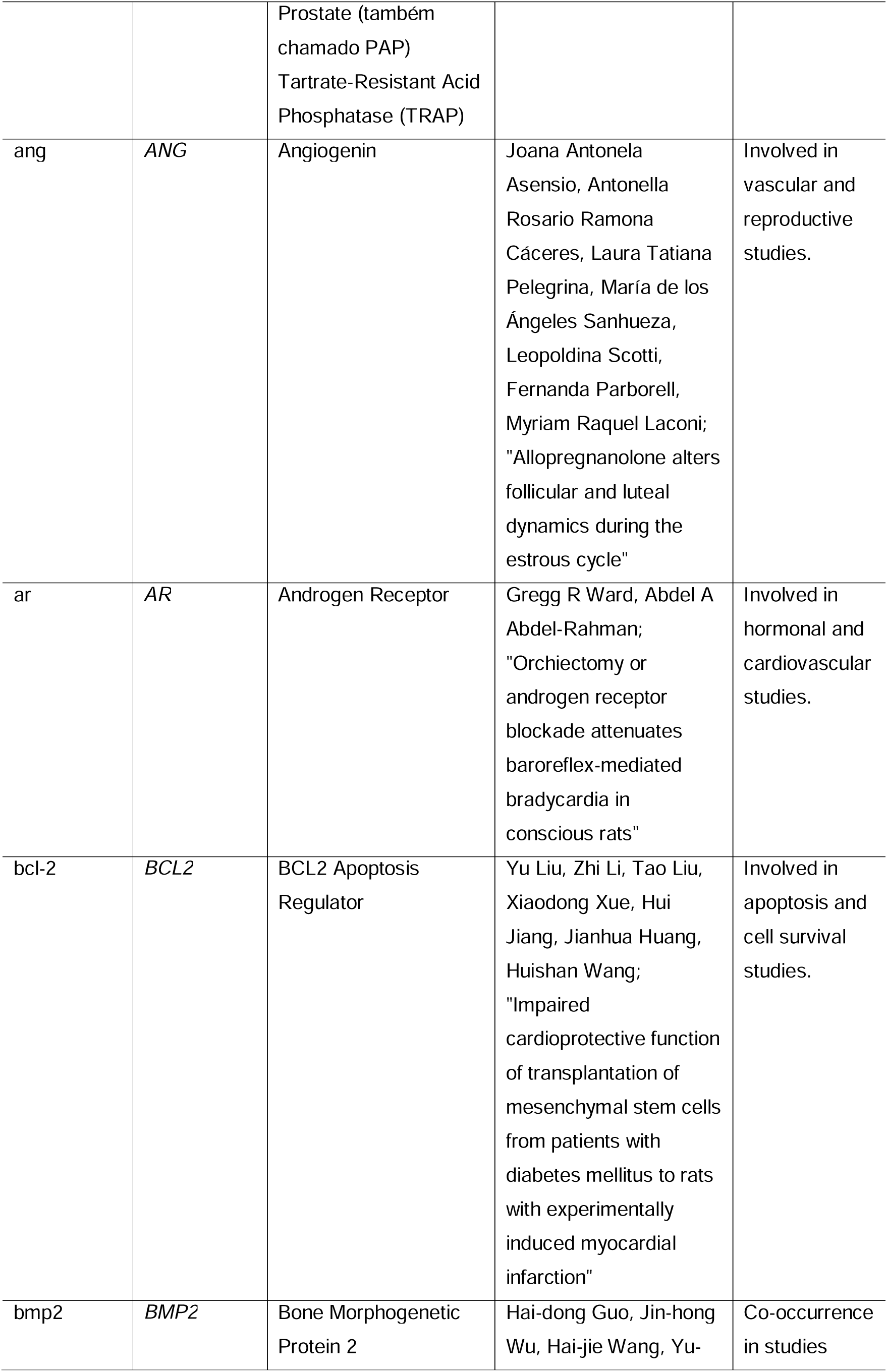
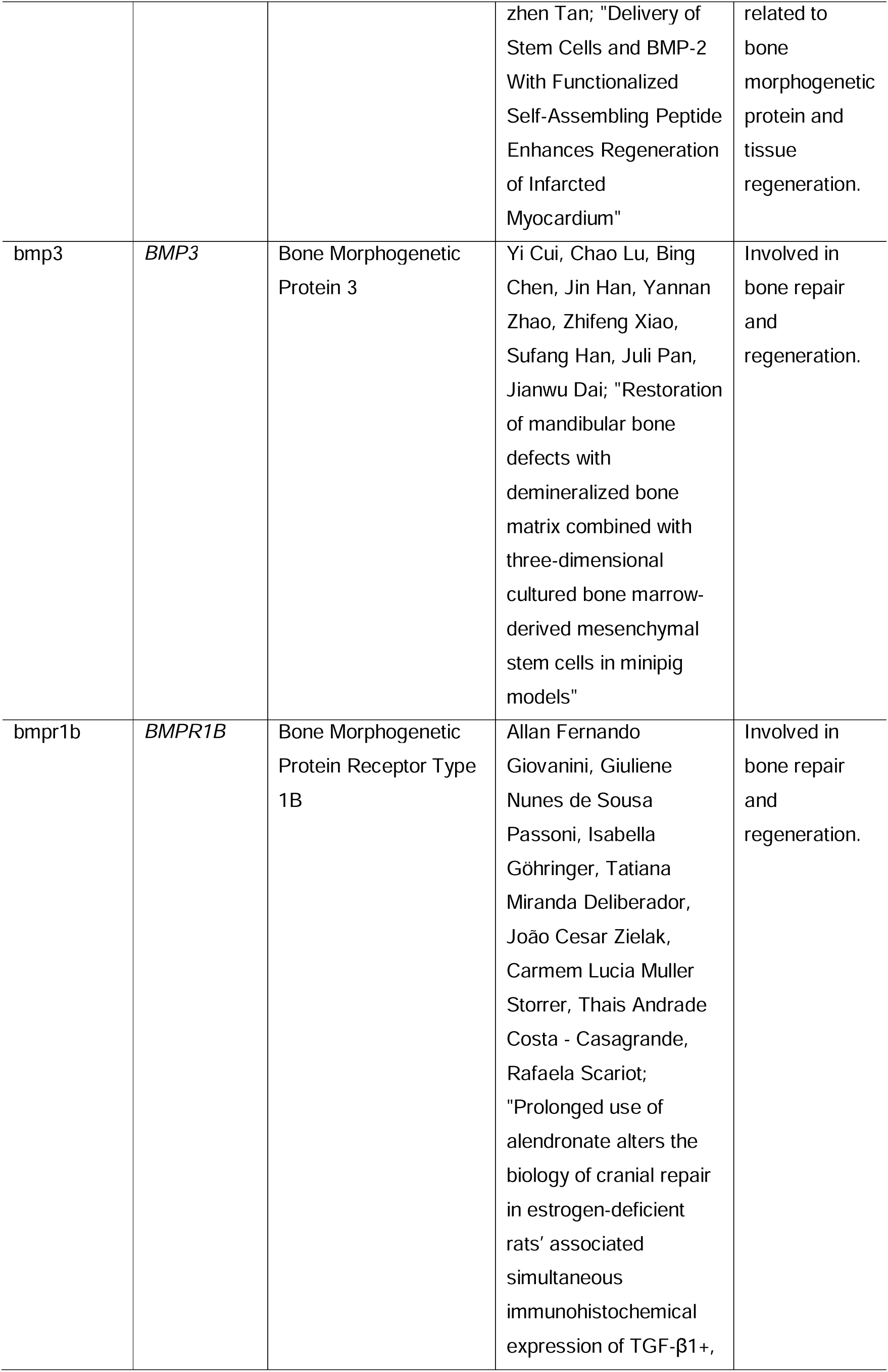
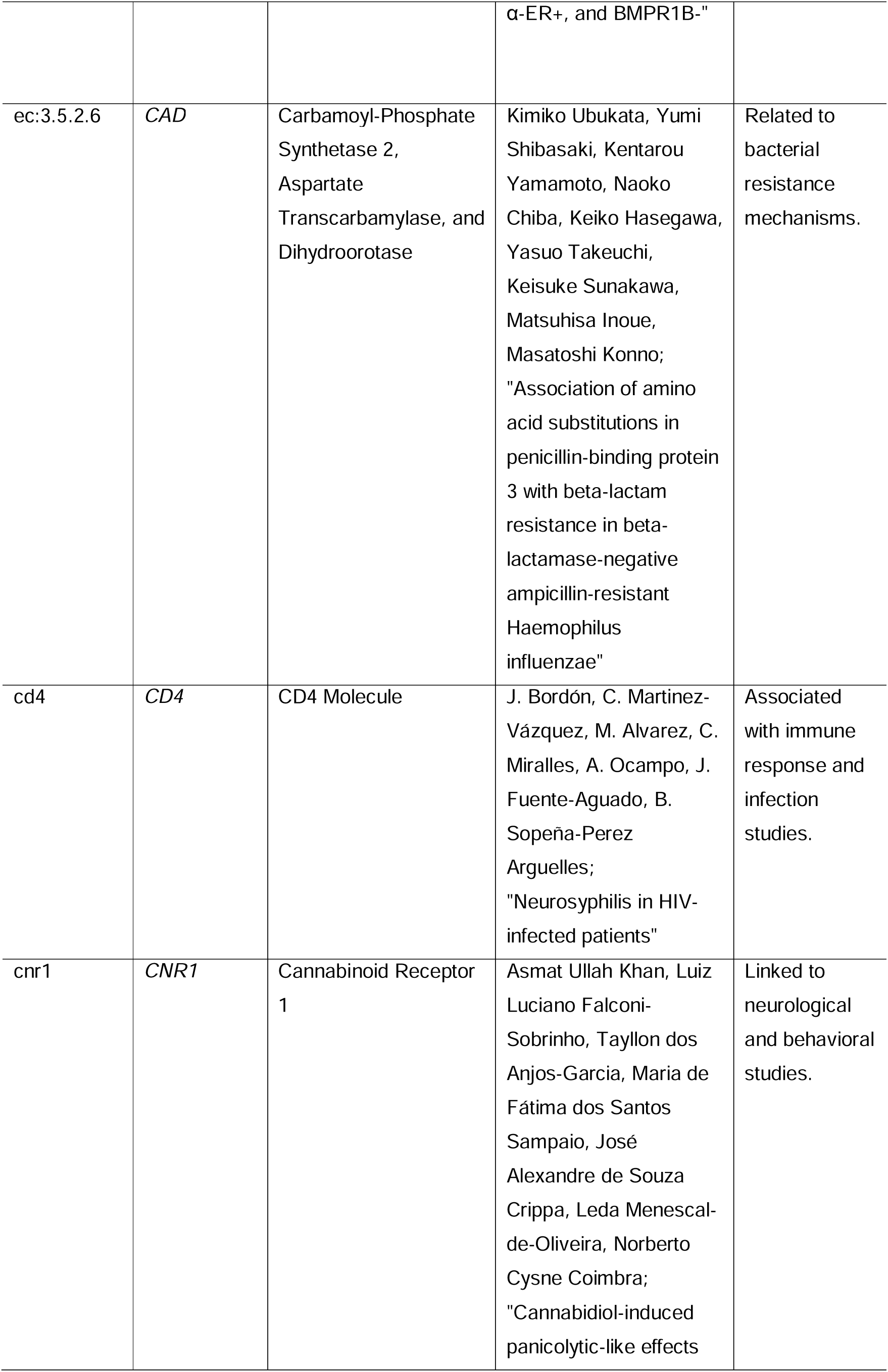
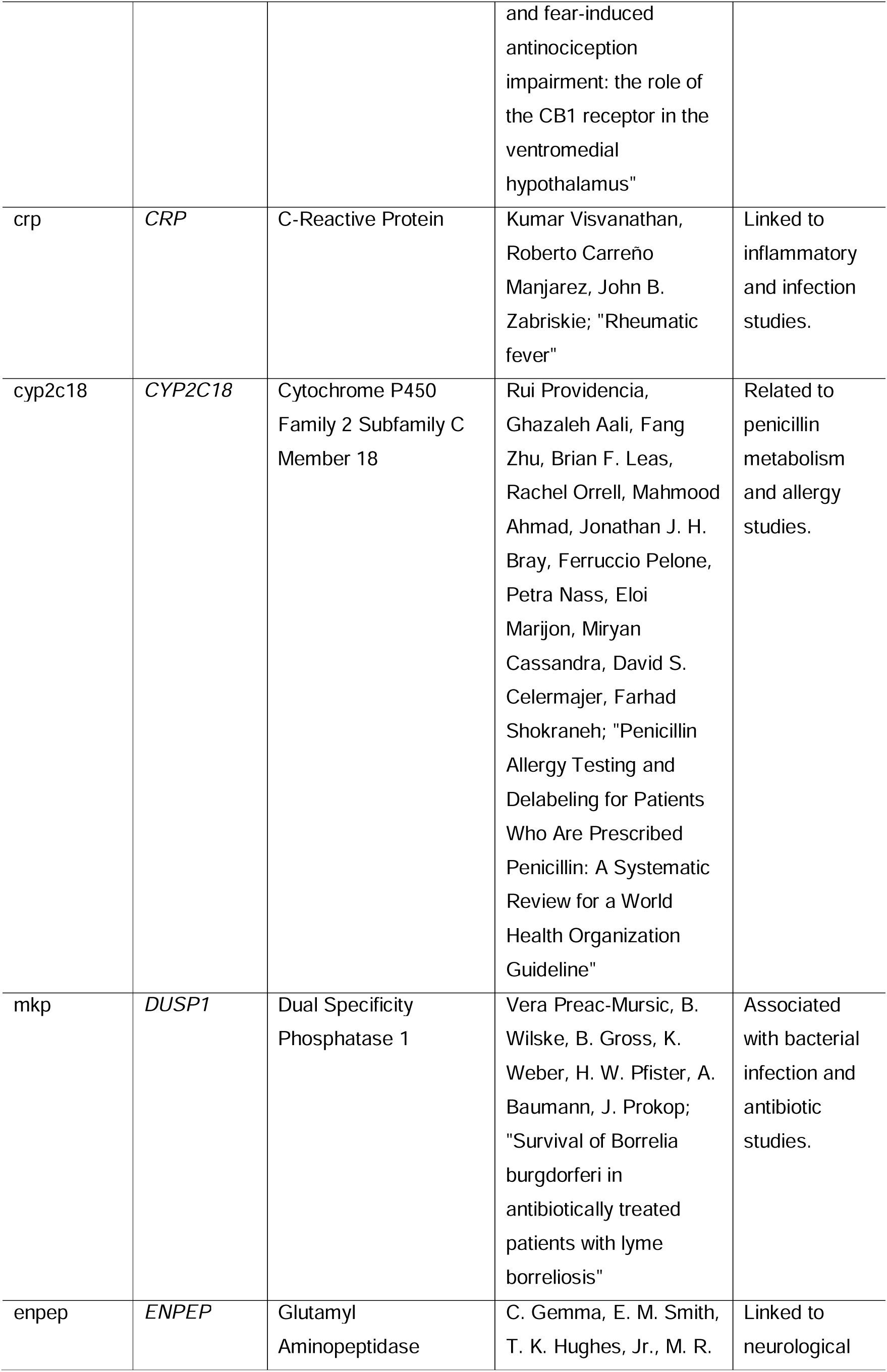
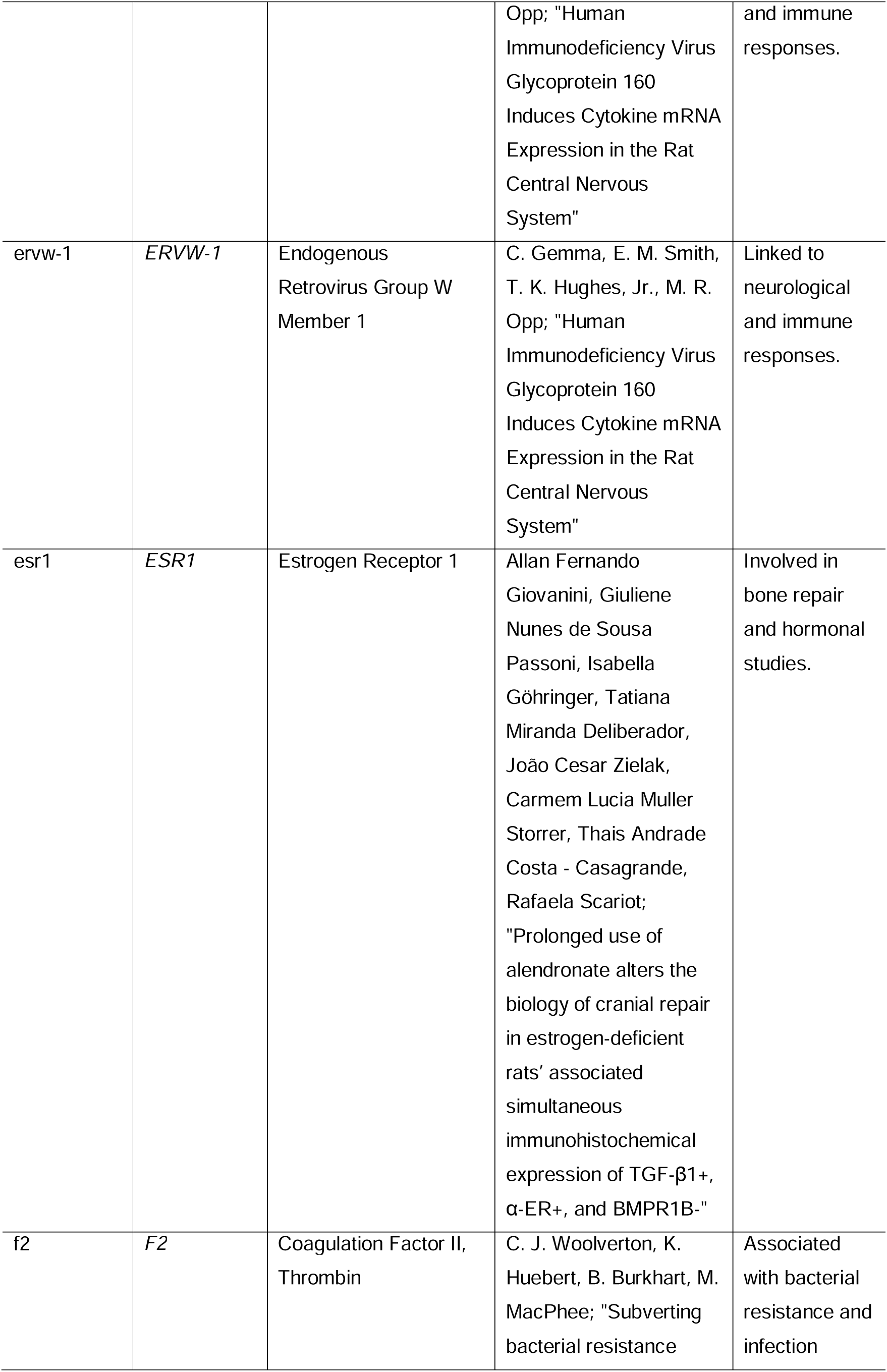
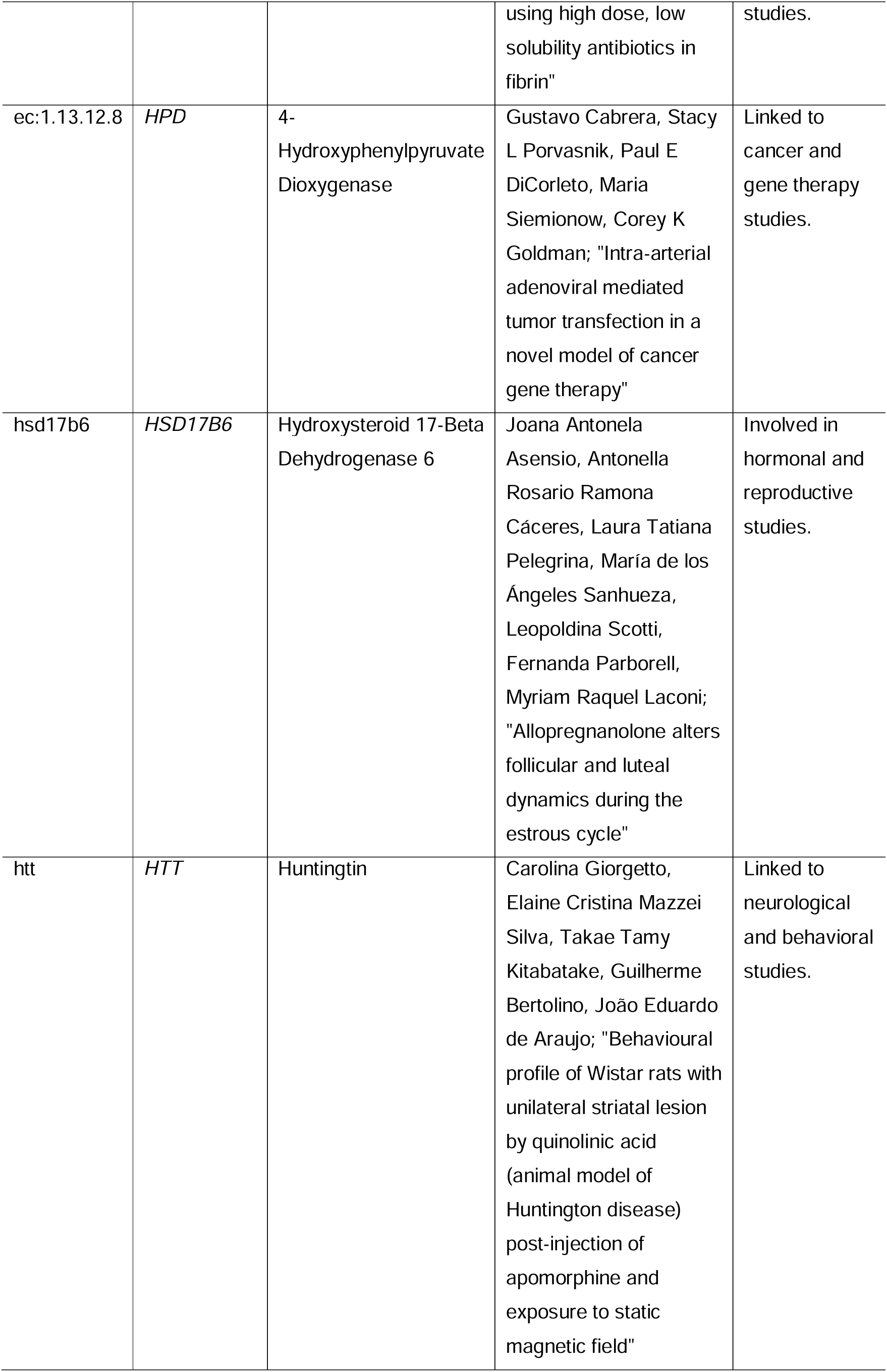
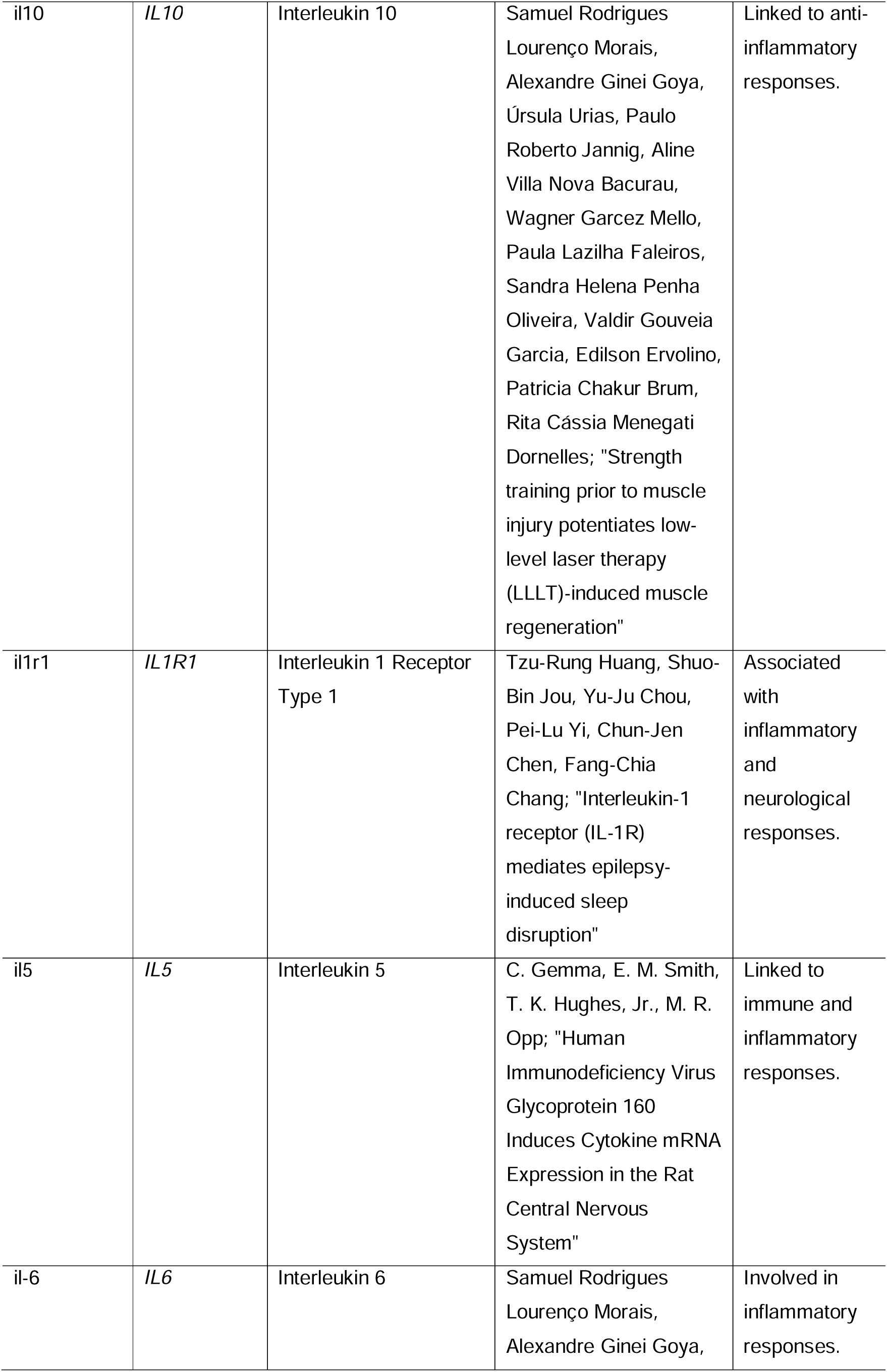
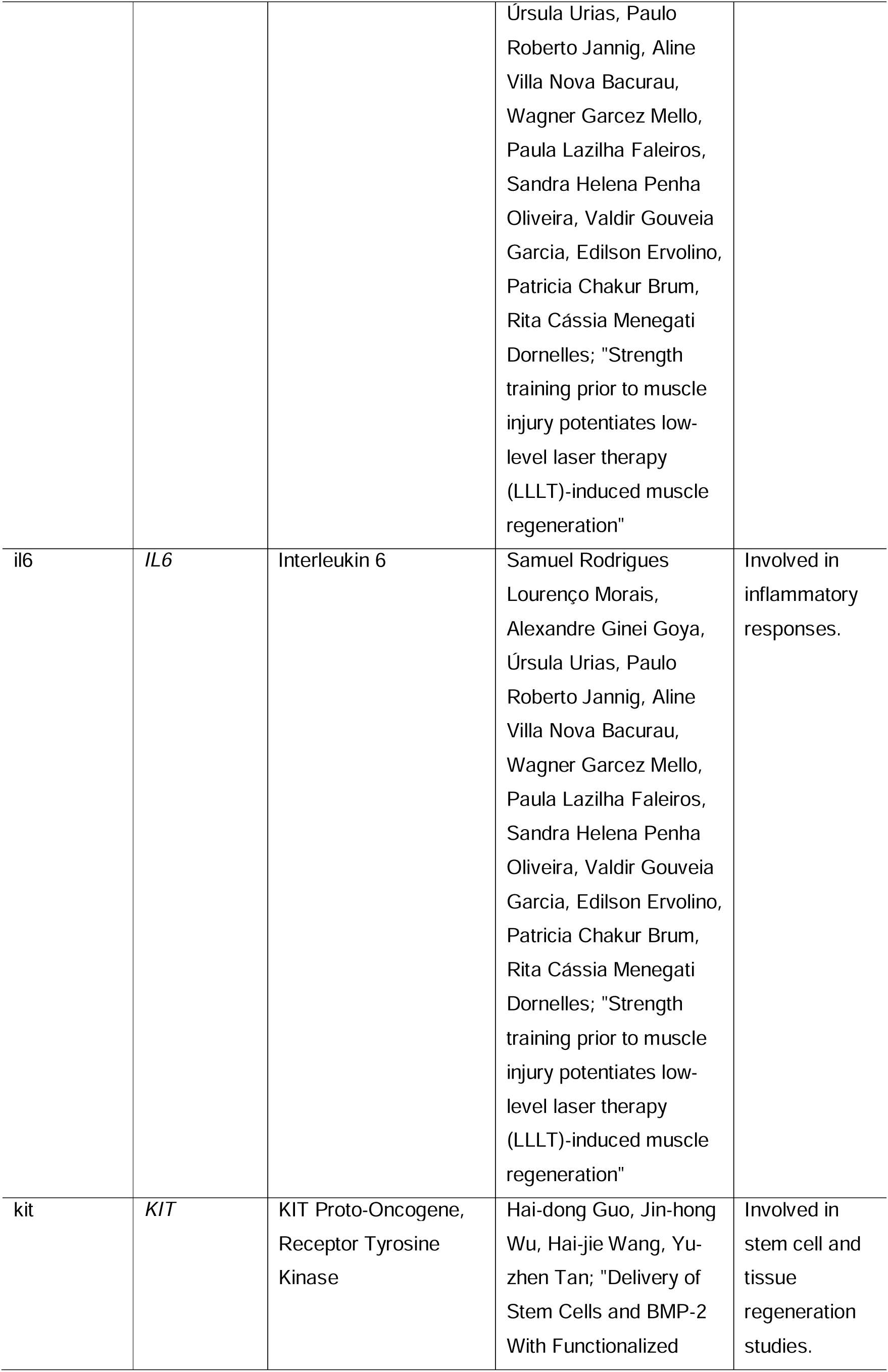
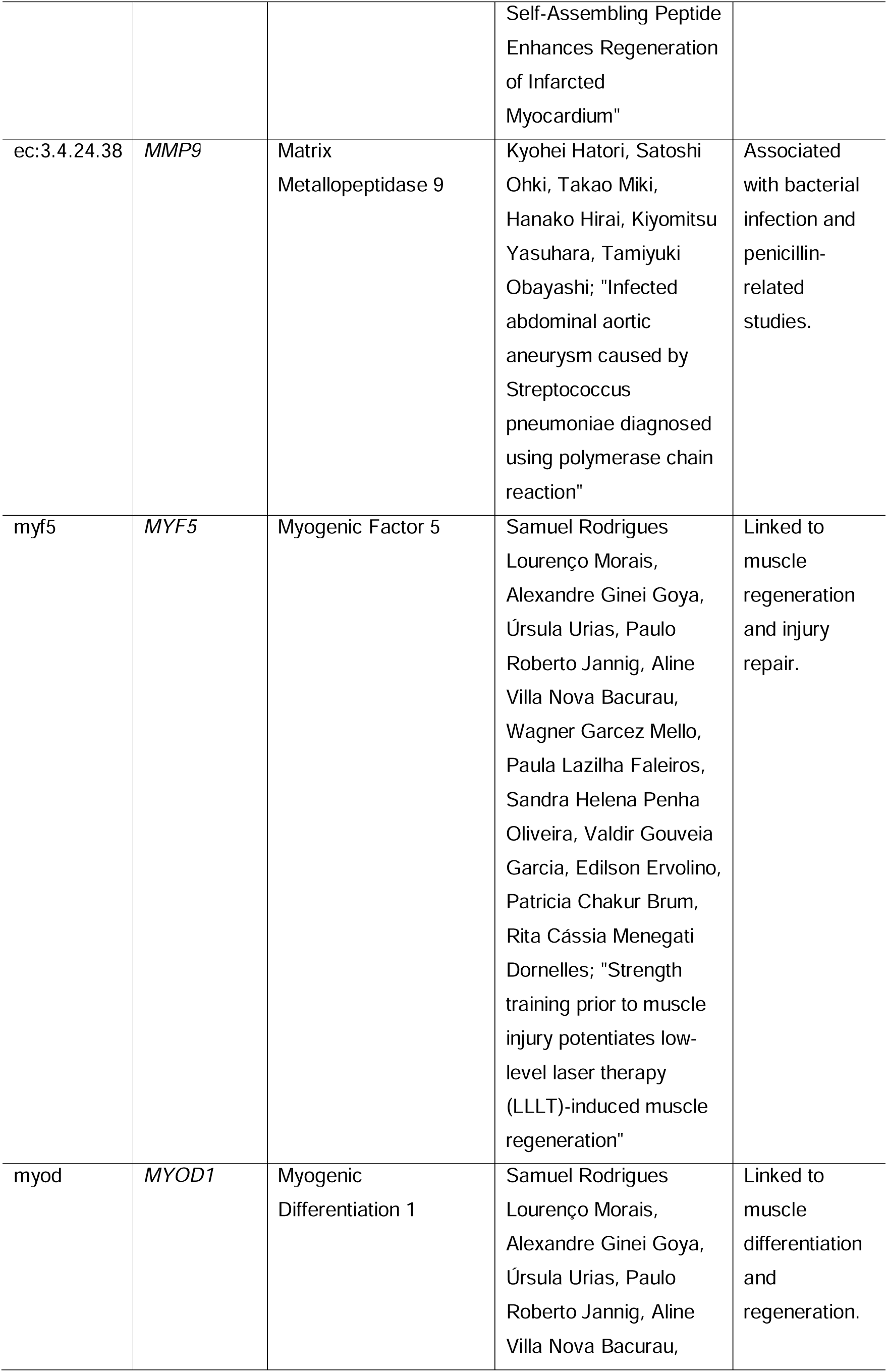
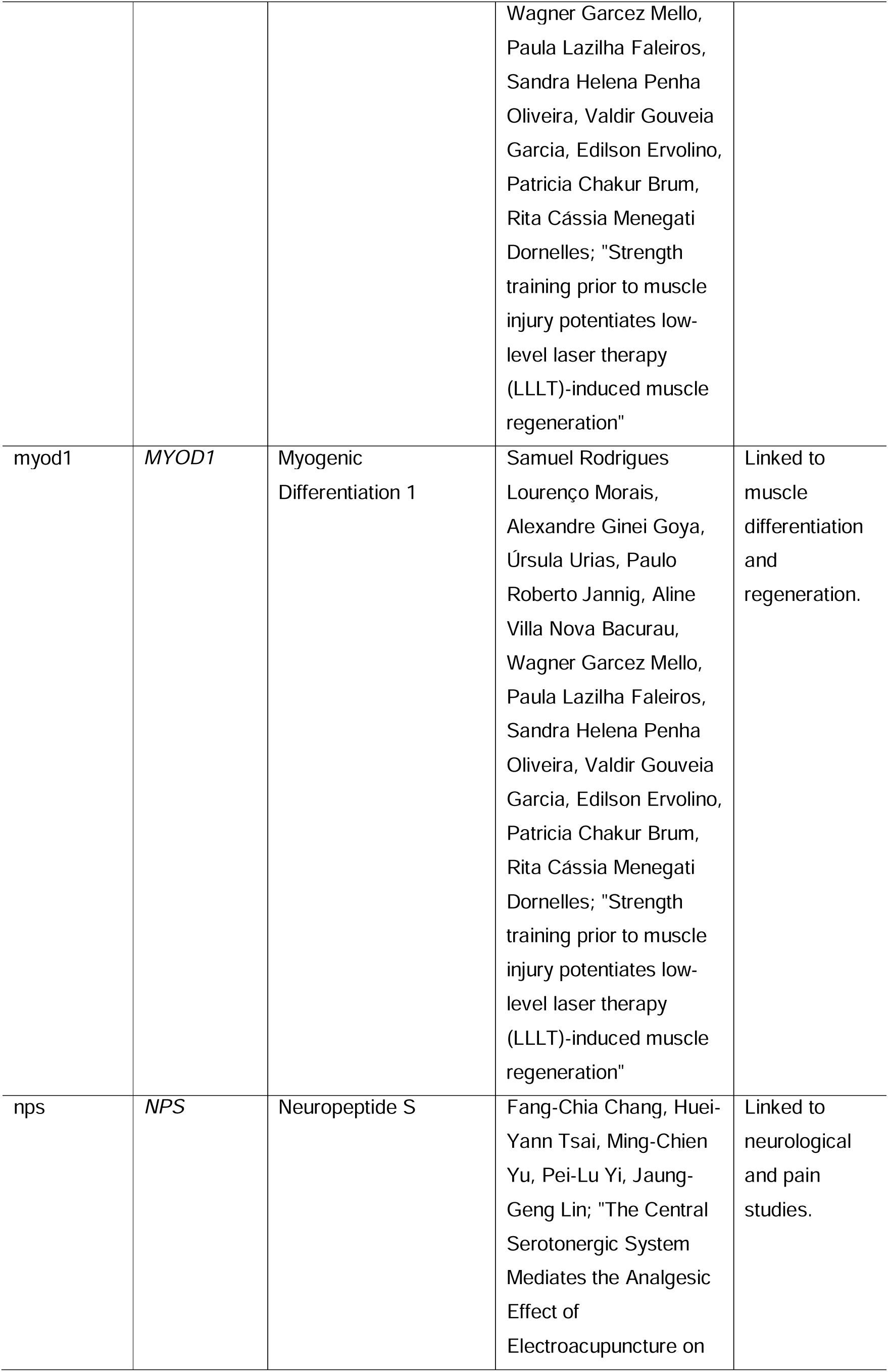
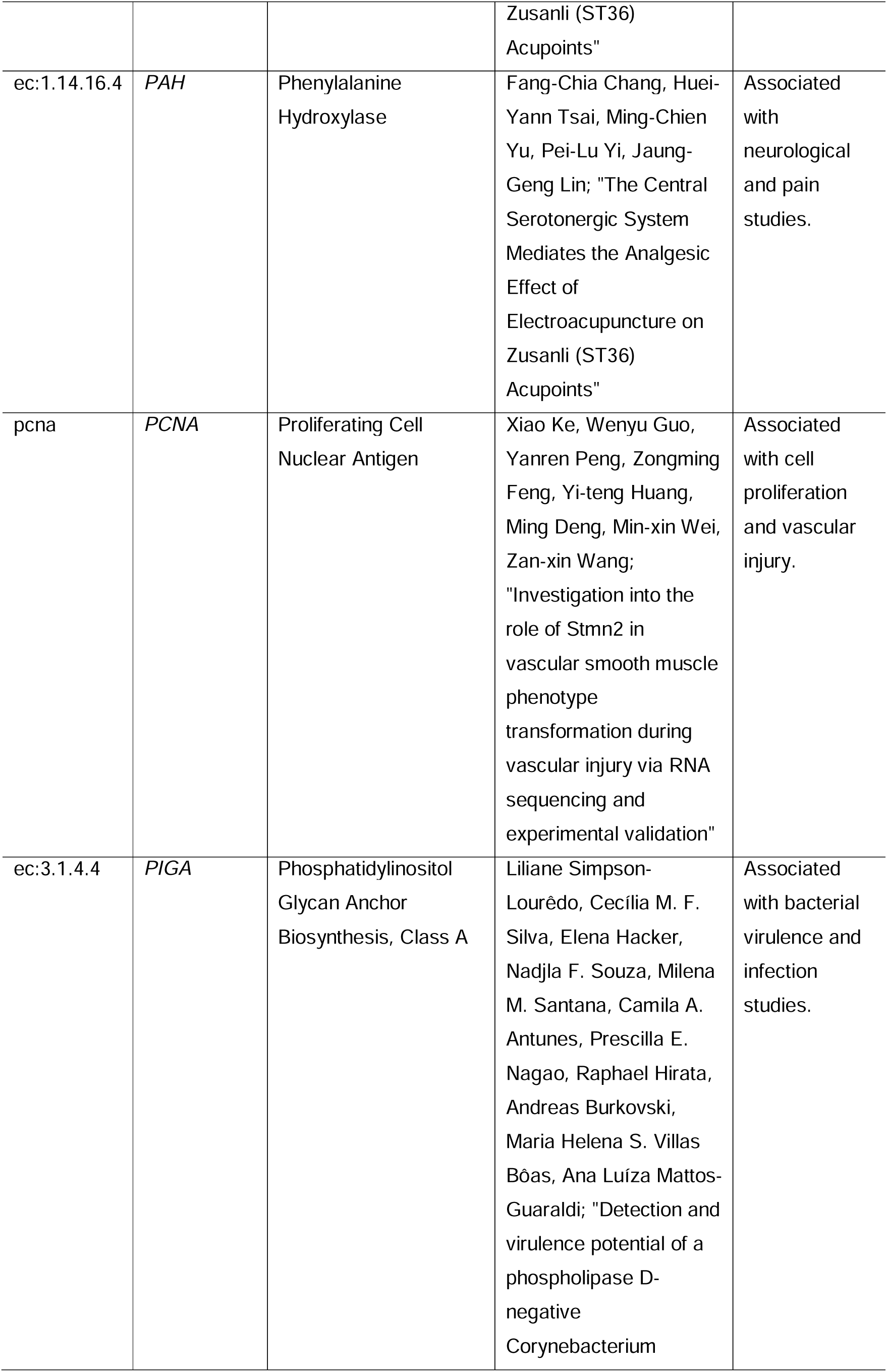
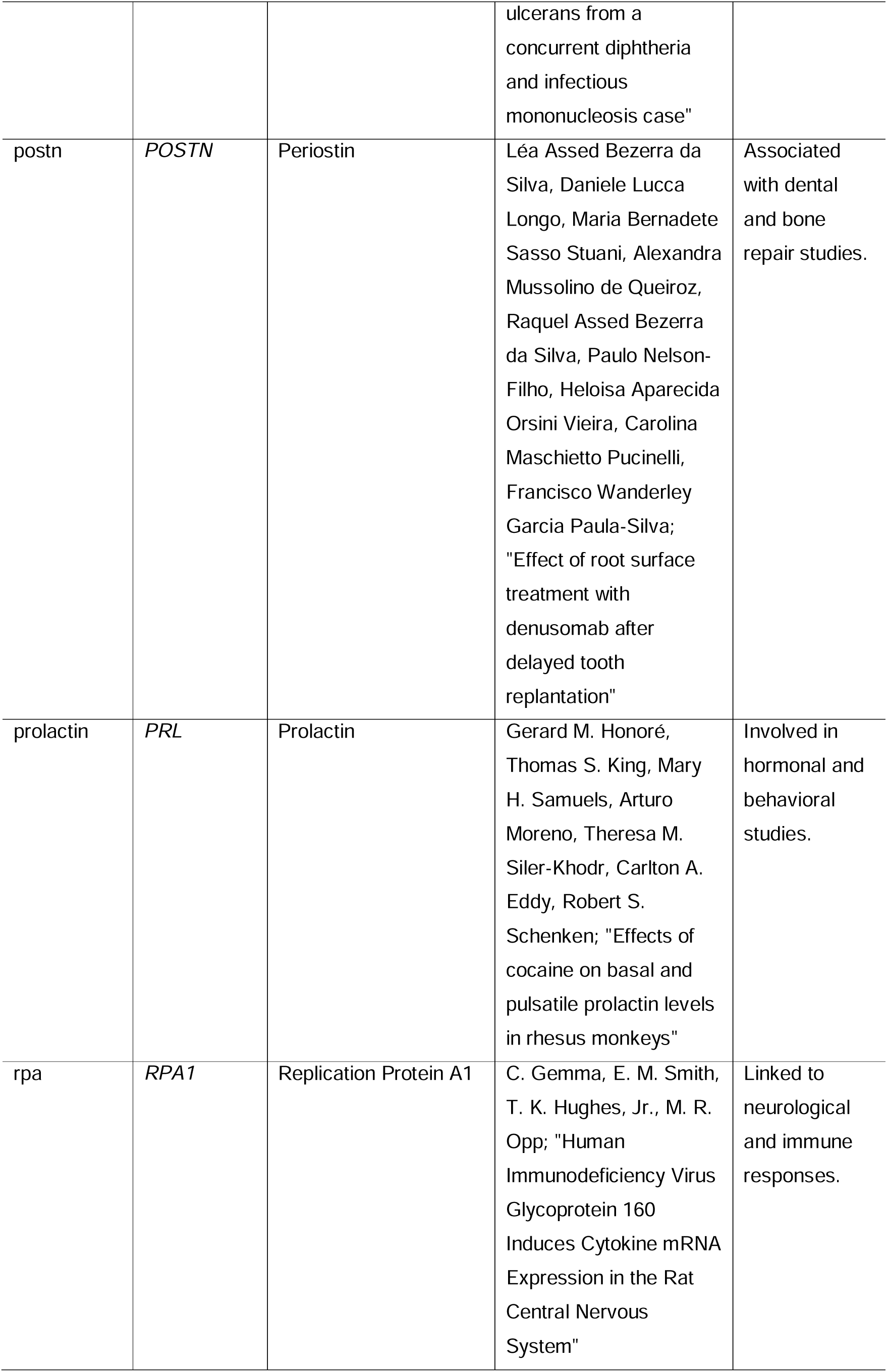
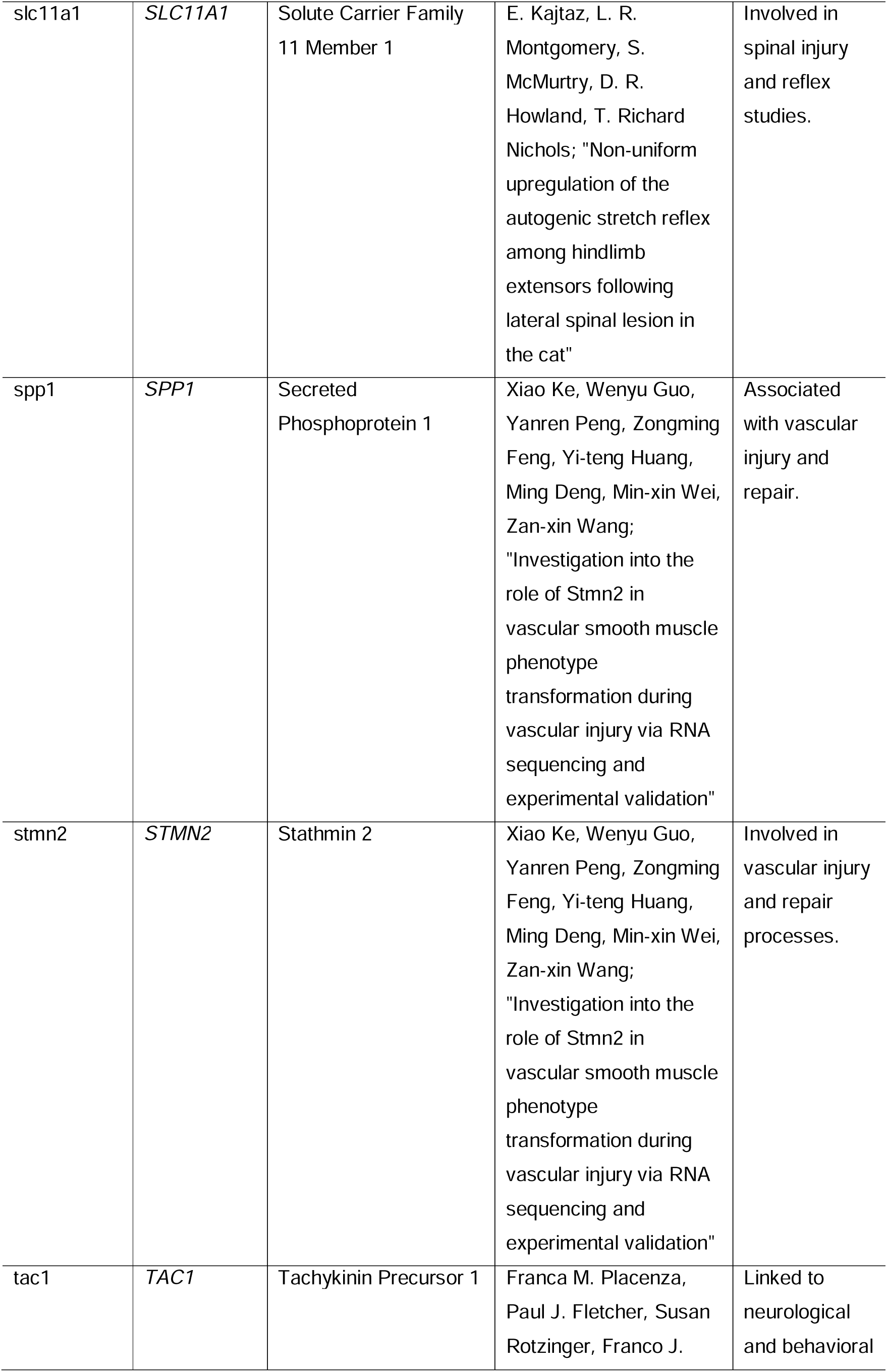
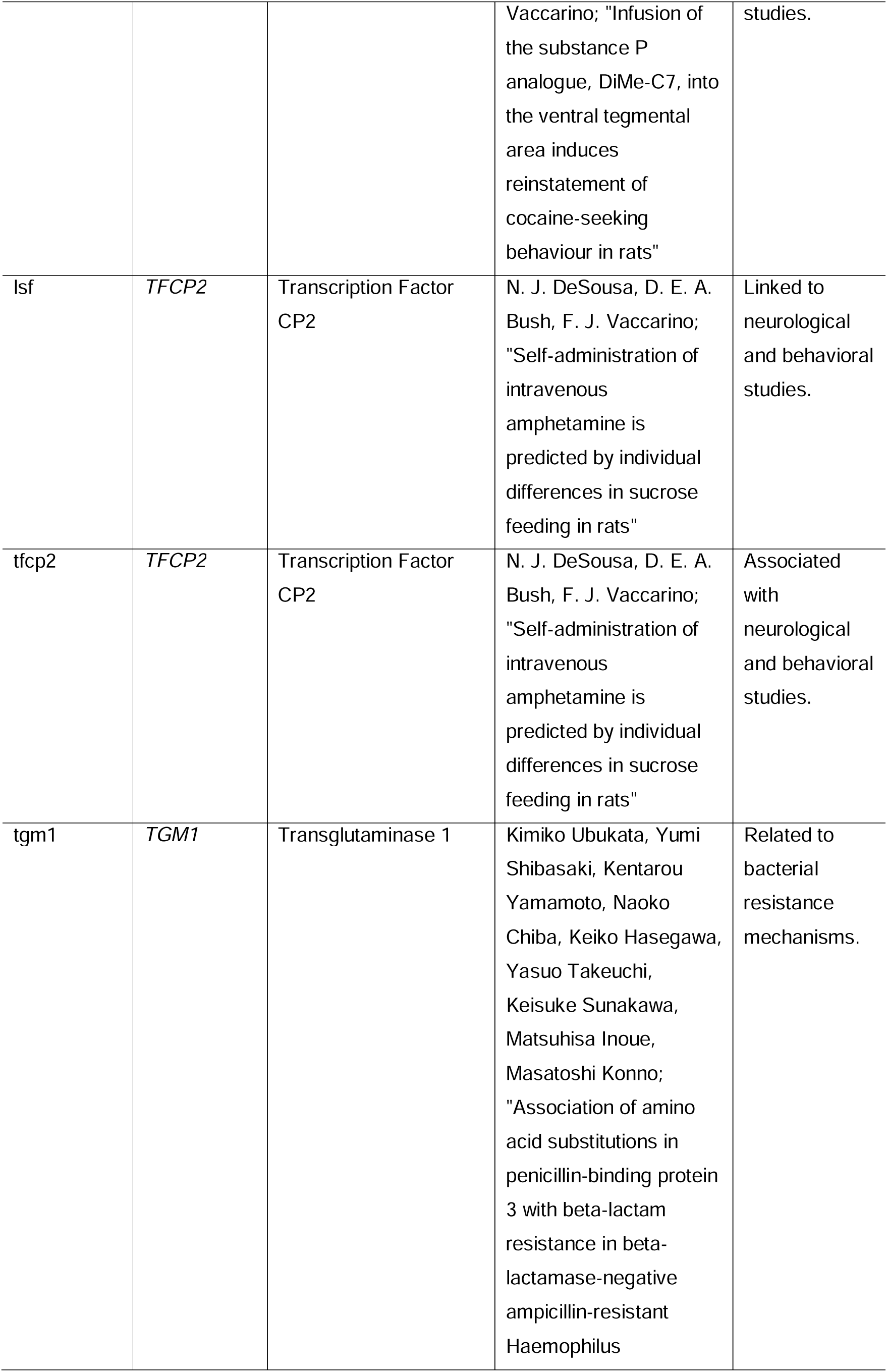
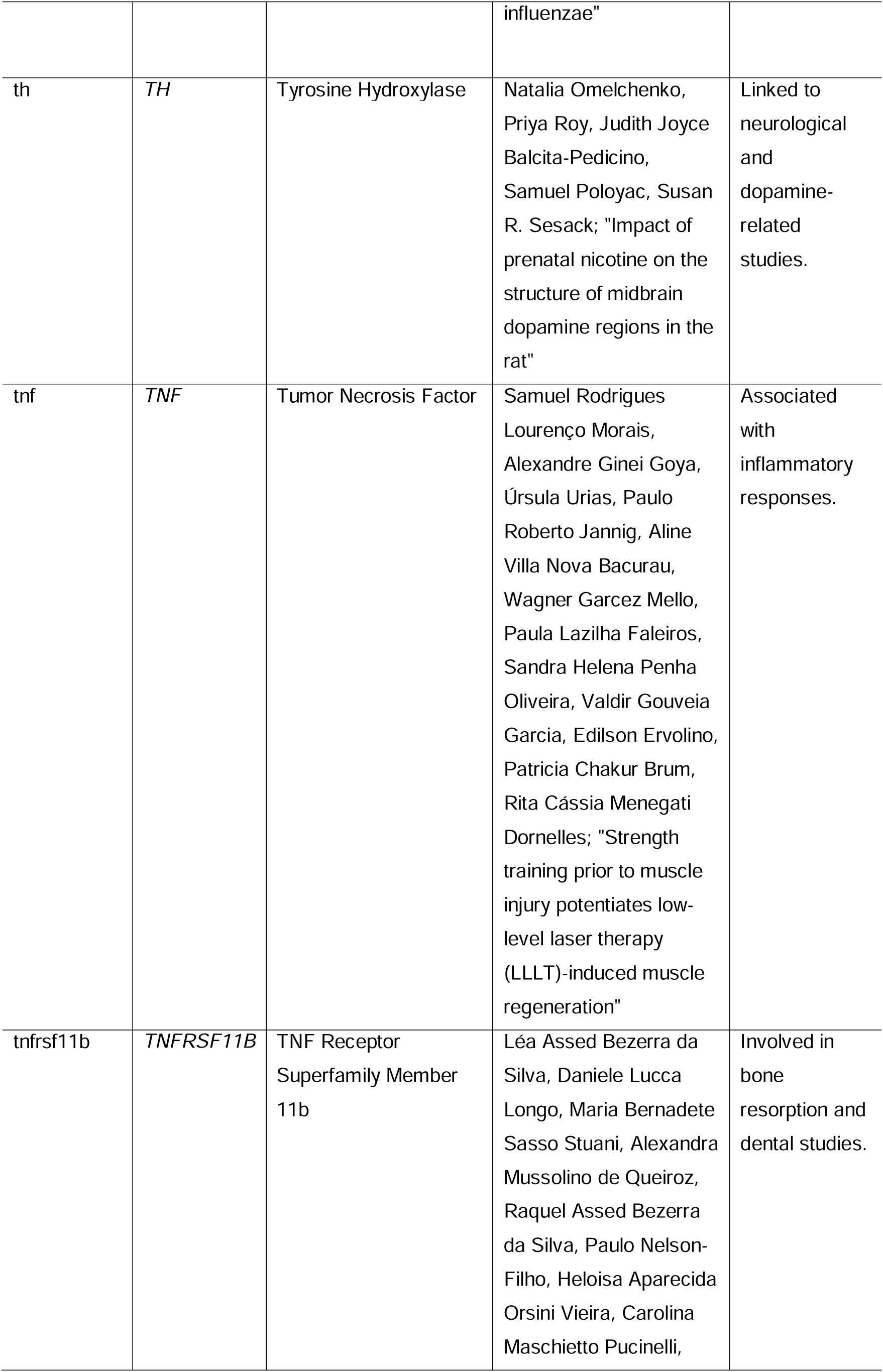
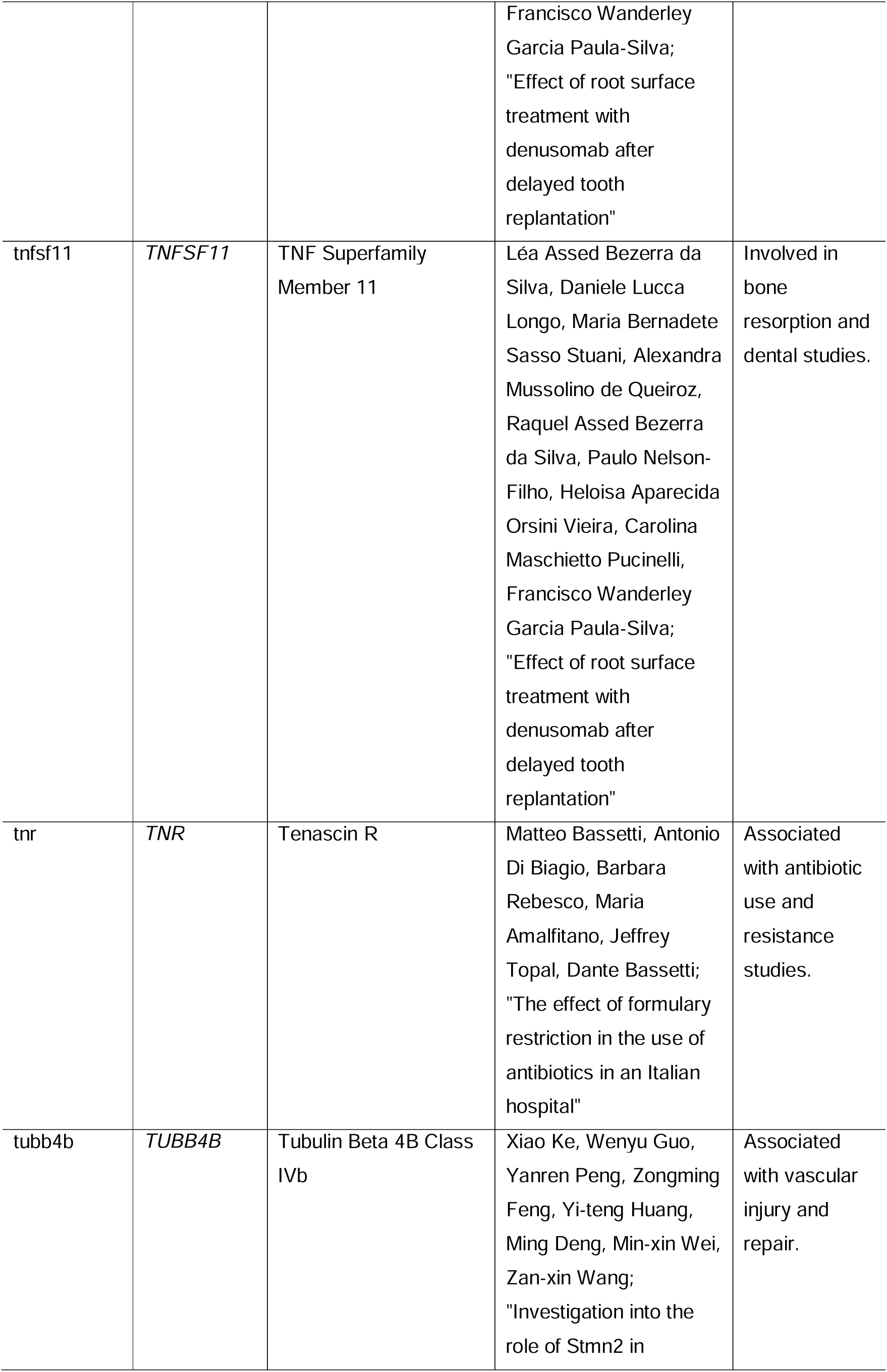
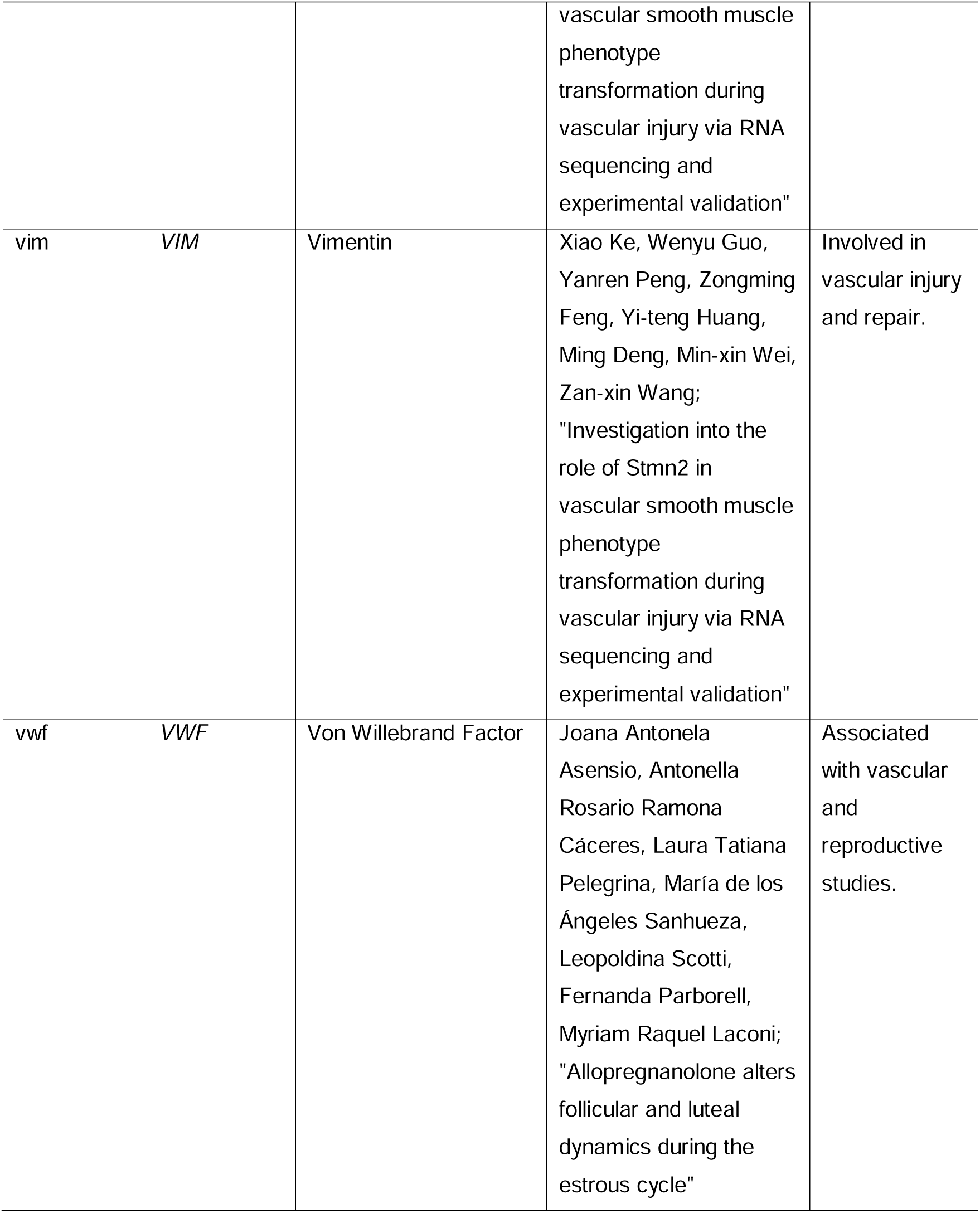
Genes associated with Penicillin G Benzathine.

**Table 2.**
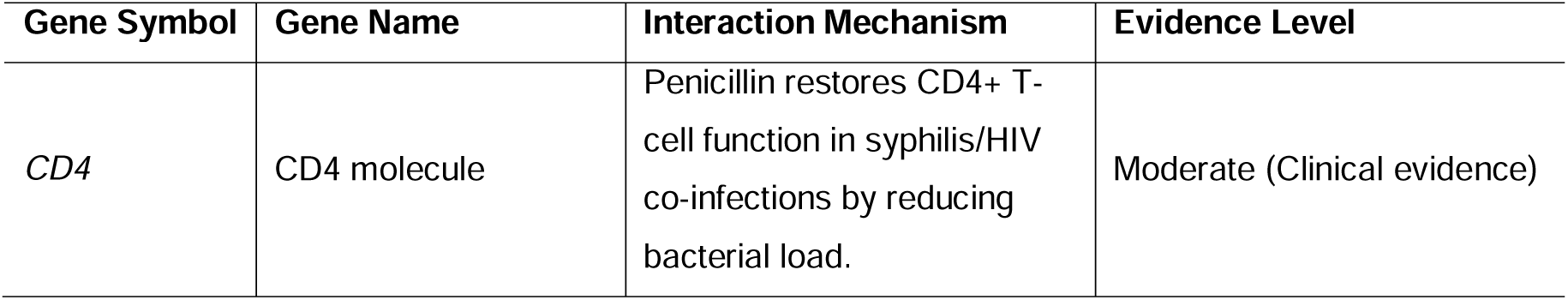
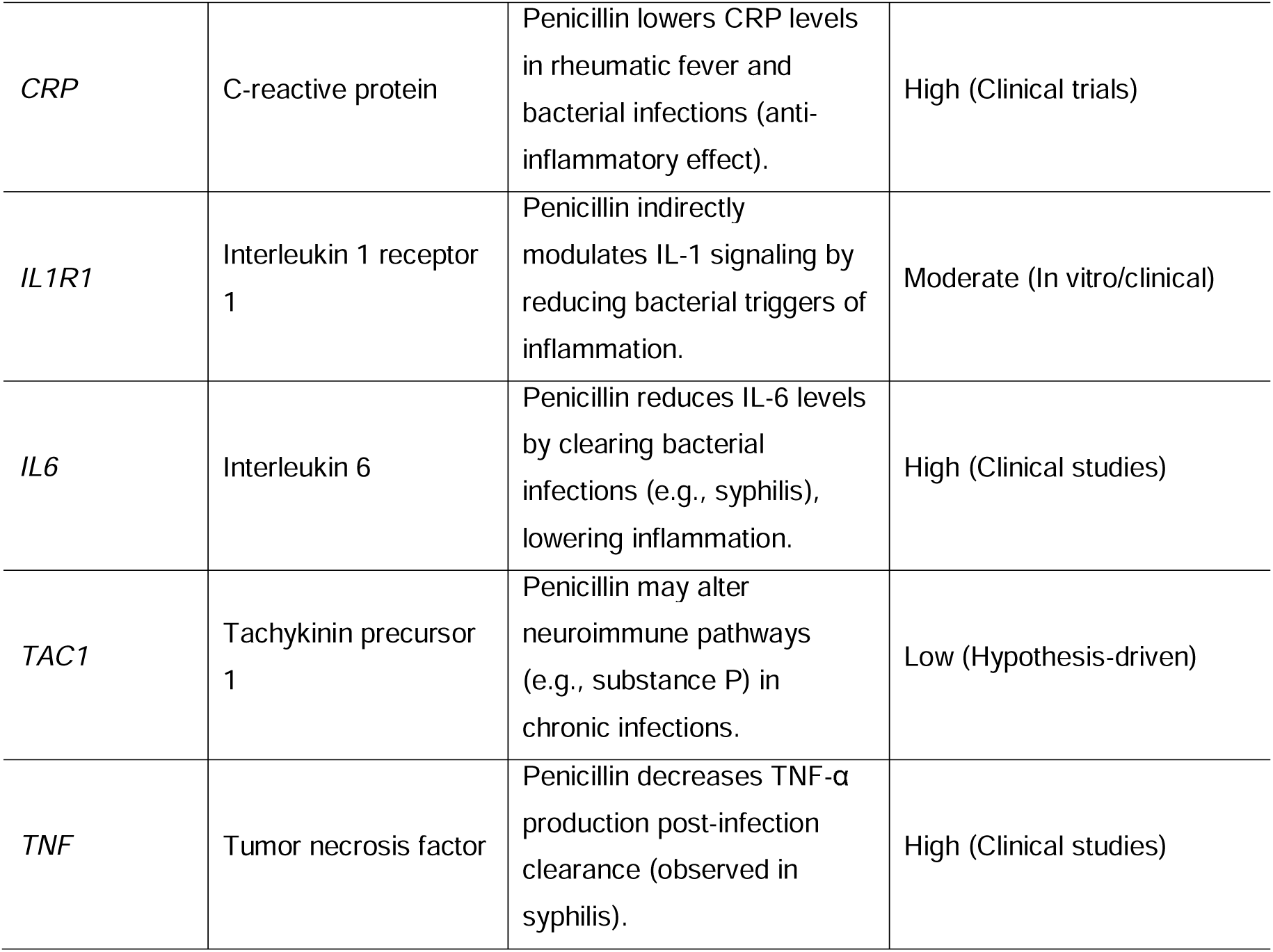
Human Genes Directly Affected by Penicillin G Benzathine.

| Gene Symbol | Gene Name | Interaction Mechanism | Evidence Level |
| --- | --- | --- | --- |
| CD4 | CD4 molecule | Penicillin restores CD4+ T-cell function in syphilis/HIV co-infections by reducing bacterial load. | Moderate (Clinical evidence) |
| *CRP* | C-reactive protein | Penicillin lowers CRP levels in rheumatic fever and bacterial infections (anti-inflammatory effect). | High (Clinical trials) |
| *IL1R1* | Interleukin 1 receptor 1 | Penicillin indirectly modulates IL-1 signaling by reducing bacterial triggers of inflammation. | Moderate (In vitro/clinical) |
| *IL6* | Interleukin 6 | Penicillin reduces IL-6 levels by clearing bacterial infections (e.g., syphilis), lowering inflammation. | High (Clinical studies) |
| *TAC1* | Tachykinin precursor 1 | Penicillin may alter neuroimmune pathways (e.g., substance P) in chronic infections. | Low (Hypothesis-driven) |
| *TNF* | Tumor necrosis factor | Penicillin decreases TNF-α production post-infection clearance (observed in syphilis). | High (Clinical studies) |

Next, we checked the expression of these 6 genes in distinct parts of the body by using the online resources GTEx Portal Database (https://www.gtexportal.org/), the Human Protein Atlas (Sjöstedt, Zhong et al. 2020) and the EYEDB for gene expression in ocular tissue (Wolf, Boneva et al. 2022). The expression analysis of the BP-modulated genes identified that the 6 genes directly modulated by BP are all expressed in several organs of the body, including in the adipose tissue, brain parts, skin, whole blood and adipose tissue (Figures S1-S6). Moreover, all genes are highly expressed in several parts of the brain (Figures 1-2), except the gene CRP, which expression was not detected in all 3 databases containing gene expression data: HPA, GTEx and FANTOM5 (Figure 1B, Figure S2). In the eye, the 6 genes directly modulated by BP are all expressed, but in different parts of the eye (Figure 3). The gene CD4 is very highly expressed in the retinal microglia and hyalocytes. IL6 is expressed more in choroid and CNV membrane, while TAC1 is expressed more in cornea and choroid.

**Figure 1.**
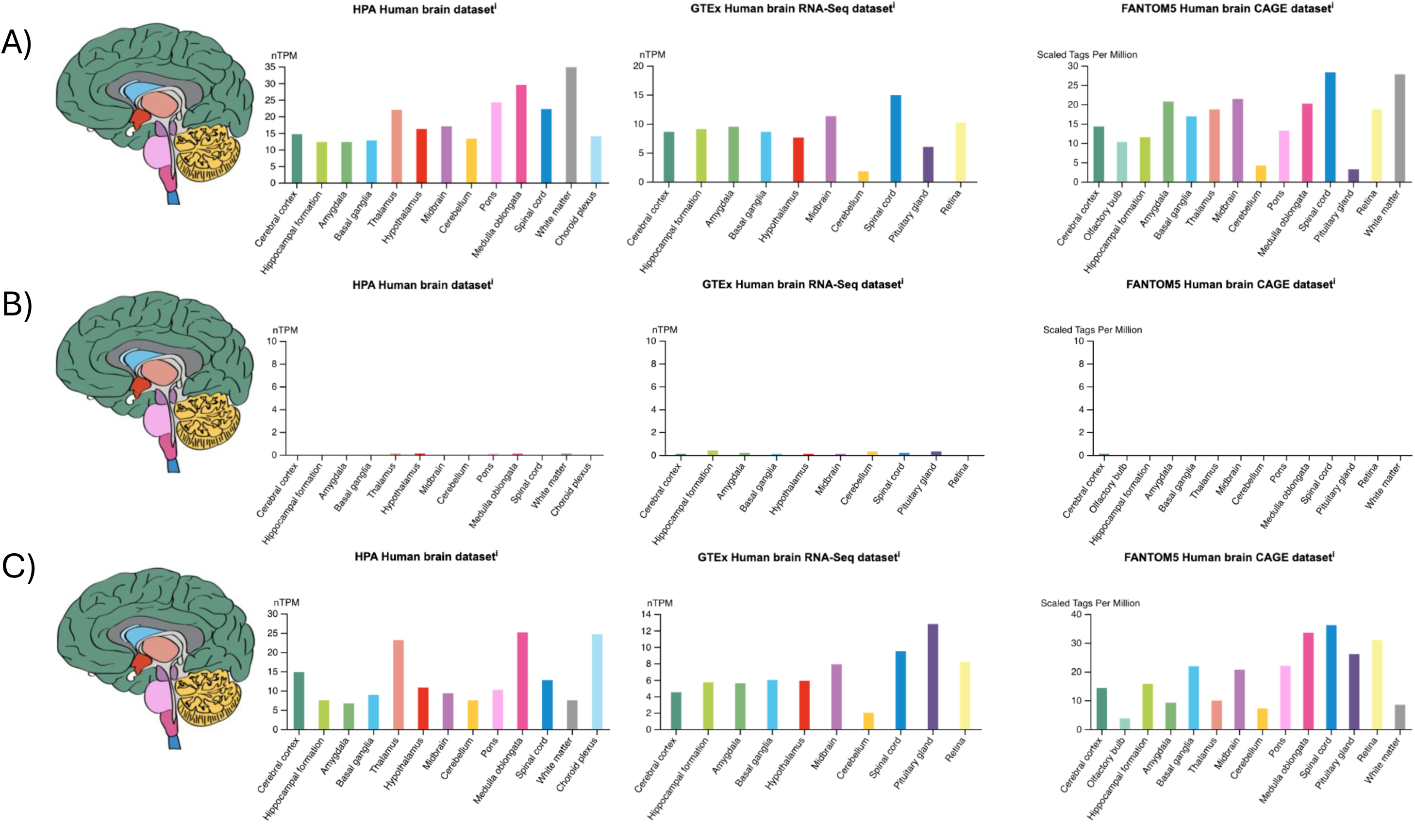
Gene expression profile of the genes CD4, CRP and IL1R1 in distinct parts of the brain in the databanks HPA, GTEx and FANTOM5. A) Expression profile of the gene CD4; B) Expression profile of the gene CRP; C) Expression profile of the gene IL1R1.

**Figure 2.**
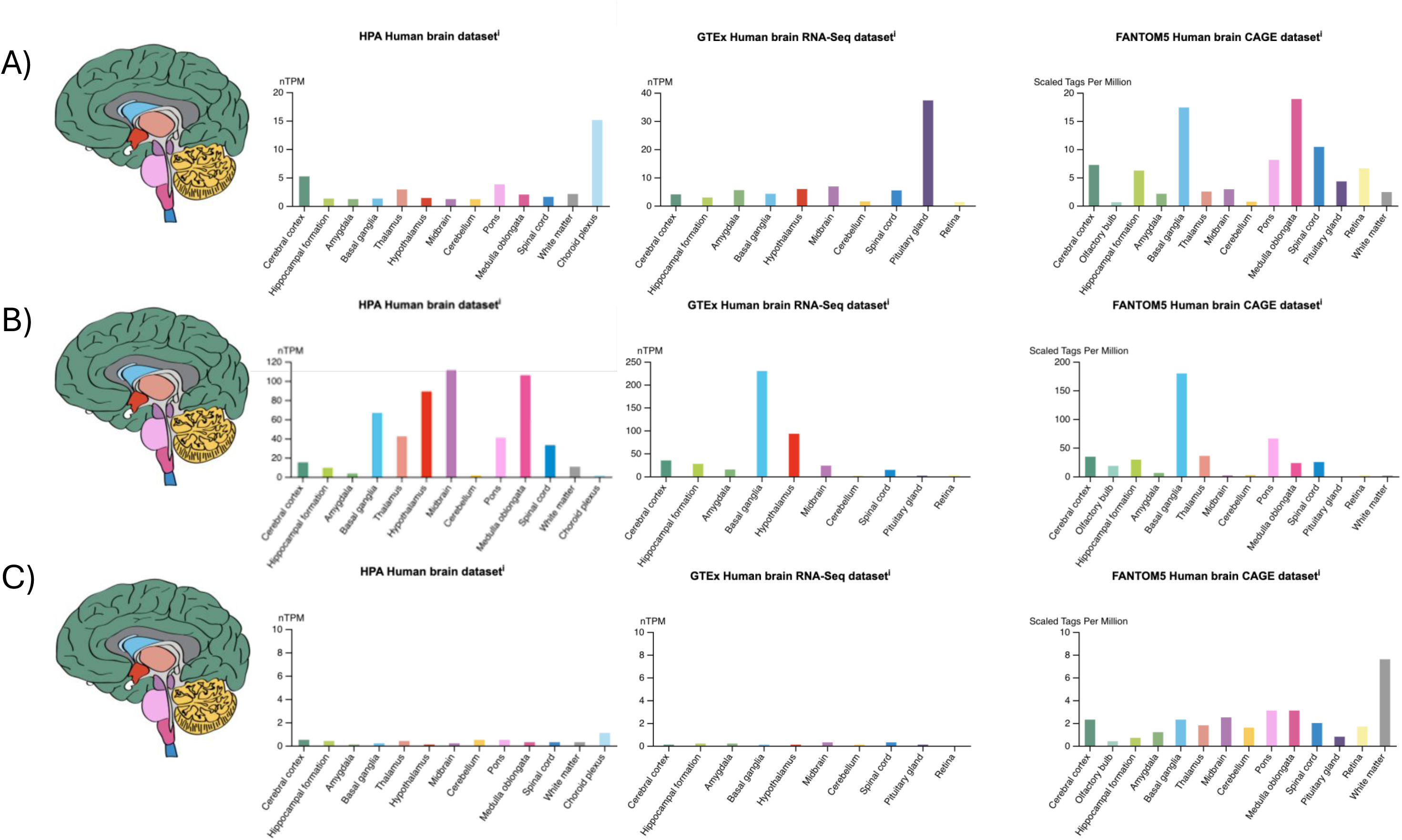
Gene expression profile of the genes IL6, TAC1 and TNF in distinct parts of the brain in the databanks HPA, GTEx and FANTOM5. A) Expression profile of the gene IL6; B) Expression profile of the gene TAC1; C) Expression profile of the gene TNF.

**Figure 3.**
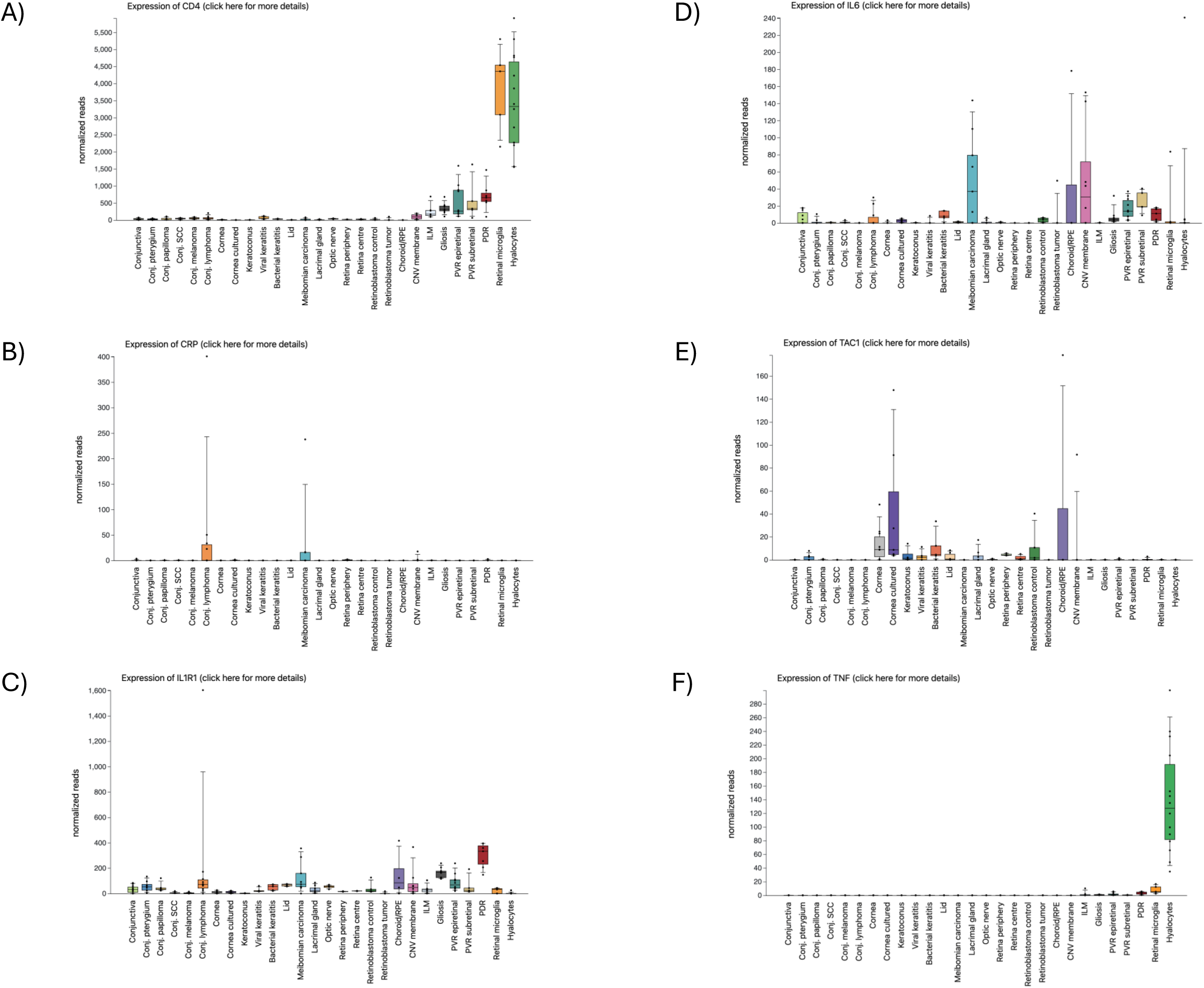
Gene expression profile of the genes CD4, CRP, IL1R1, IL6, TAC1 and TNF in eye parts. A) Expression profile of the gene CD4 in the eye; B) Expression profile of the gene CRP in the eye; C) Expression profile of the gene IL1R1 in the eye; D) Expression profile of the gene IL6 in the eye; E) Expression profile of the gene TAC1 in the eye; F) Expression profile of the gene TNF in the eye.

According to the literature, these genes are involved in several biological processes that play an important role in eye development and/or maintenance. The gene *TAC1* (Tachykinin Precursor 1) is a coding gene related to pre-pro-peptide receptors called pre-pro-tachykinin A, which undergoes post-translational cleavages to generate several active peptides, including: substance P, neurokinin A (NKA), and NKA variants (NPK and NPγ) (Peng, Agogo et al. 2019). In the retina, substance P acts in the recovery part of RPE cells injured by oxidative stress. This mechanism is associated with macular degeneration (Baek, Yu et al. 2016).

The gene *IL1R1* (Interleukin 1 Receptor Type 1) encodes a receptor of the interleukin-1 family that interacts with: interleukin-1 alpha, IL-1beta and interleukin-1 receptor agonist (Mosley, Urdal et al. 1987). IL1beta and IL1R1 signaling are associated as a protective factor for retina through microglia in an animal model (Todd, Palazzo et al. 2019).

The gene *CRP* (C-Reactive Protein) codes for a protein associated with recognizing pathogens and damaged host cells and initiating their elimination (Kilpatrick and Volanakis 1991). In the pathways of inflammation and complement activation, it is associated with other genes described in the *IL6* and *TNF* that increase its expression (Blake and Ridker 2002). AMD (AMD) is associated with higher expression of CRP (Subhi, Krogh Nielsen et al. 2019) and its pathogenesis (Chirco and Potempa 2018).

The gene *CD4* (CD4 Molecule) encodes the membrane glycoprotein of T lymphocytes (Zeitlmann, Sirim et al. 2001). It is associated with the pathogenesis of AMD through inflammation and complement pathways (Zeng, Yin et al. 2021). *CFH* gene polymorphisms are associated with a decrease in CD4+ cells and associated with AMD pathogenesis (Nielsen, Subhi et al. 2023).

The gene *TNF* (Tumor Necrosis Factor) gene encodes a cytokine (TNF-α) that is associated with inflammation and the immune response. (Falvo, Tsytsykova et al. 2010). In the retina, TNF-α decreases the expression of genes such as CDH1, RP65, RDH5, RDH10, TYR, MERTK, MITF, TRPM1, TRPM3, and miRNAs 206 and 211 related to phagocytosis, visual cycle, and epithelial morphology and is associated with phagocytosis, visual cycle, and epithelial morphology. Prolonged exposure to TNF-α alters the retinal blood-barrier of RPE, decreases the expression of visual cycle genes such as *OTX2*, and decreases space immunosuppression. These modifications are very similar to alterations found in AMD (Touhami, Beguier et al. 2018).

The gene *IL-6* (Interleukin 6) translates to an inflammation-related cytokine (IL-6) (Schmidt-Arras and Rose-John 2016). In the retina, prolonged exposure to pro-inflammatory factors such as IL-6 and TNF-α decreases the viability of RPE and alters the blood-retinal barrier and cellular function, contributing to the development of atrophic AMD (Klettner, Brinkmann et al. 2020). Studies in animal models have revealed that IL-6 prolongs the survival of subretinal macrophages by generating chronic inflammation, related to the pathogenesis of AMD (Xiao, Lei et al. 2023). The expression of IL-6 in the retina also increases after exposure to UV light and activation of the complement system occurs, serving as a model for the study of AMD (Lueck, Hennig et al. 2012).

We also found that 4 out of these 6 genes (*IL6*, *TNF*, *IL1R1* and *TAC1*) (Table 3) that are directly modulated by BP are also widely associated with several ocular pathologies. The overexpression of IL6, for example, promotes chronic inflammation, angiogenesis, and retinal damage, leading to vision loss via edema and neovascularization. The eye diseases associated with misregulation in IL6 gene are uveitis, diabetic retinopathy and age-related macular degeneration (AMD). Another important gene is *TAC1*, and its overexpression increases alterations in cornealsensory innervations and inflammation. The eye diseases associated with misregulation in *TAC1* gene are neuropathic corneal pain and dry eye.

**Table 3.**
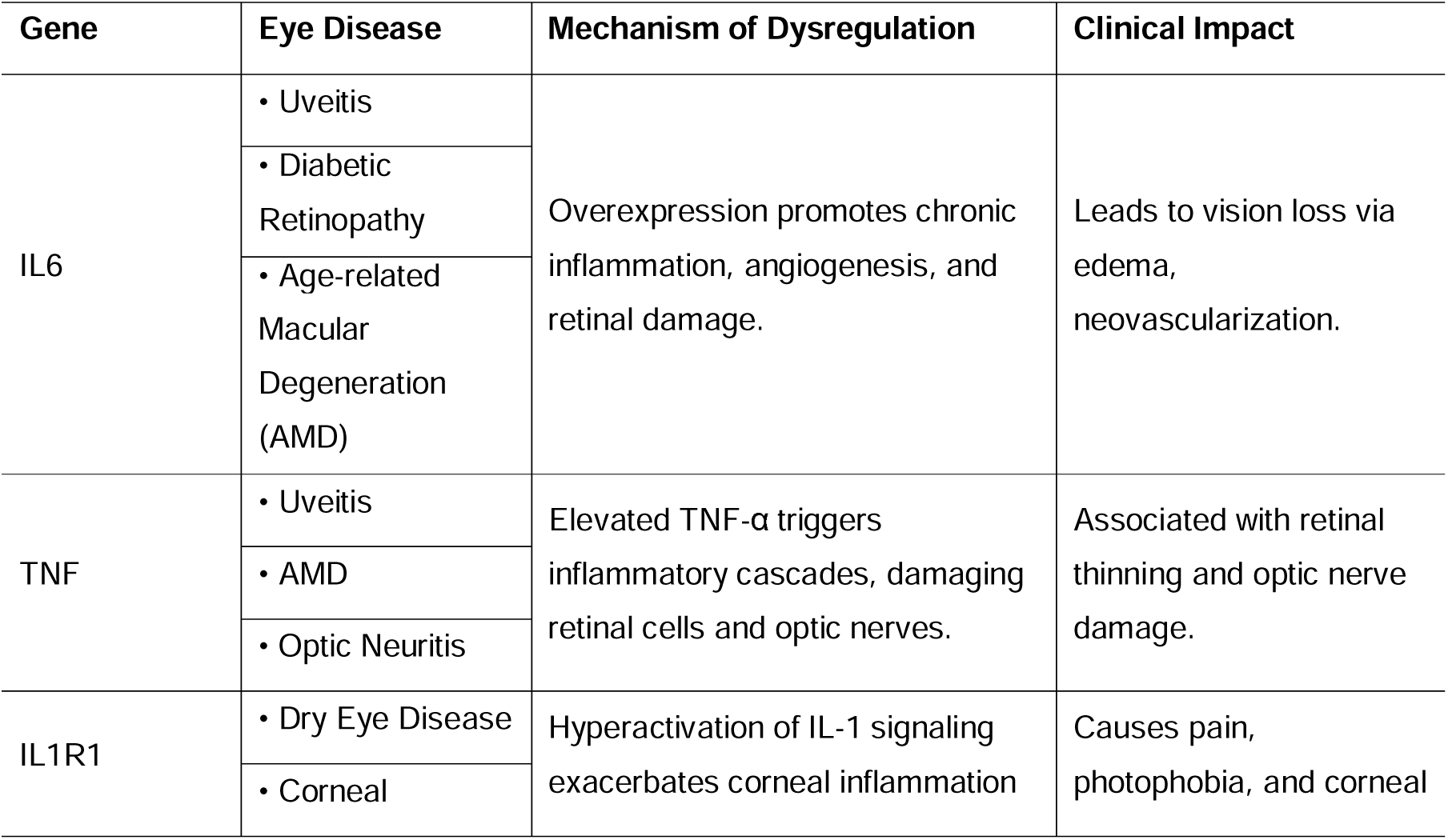
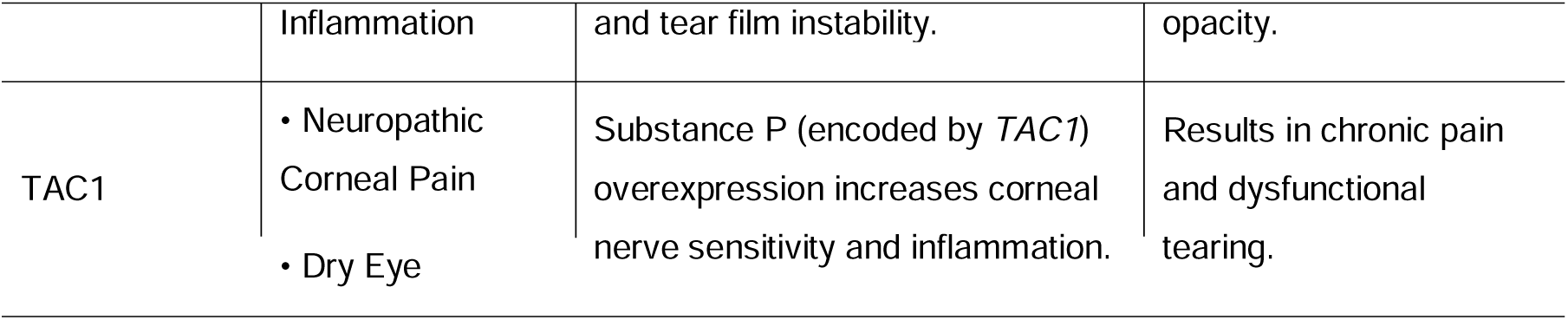
Genes Linked to Eye Diseases When Dysregulated.

Next, to investigate whether the genes modulated by BP are interacting to each other, we subjected the 6 genes to STRING software (Szklarczyk, Kirsch et al. 2023) to investigate the interaction between their coded proteins. Using a set of non-restrictive parameters (types of interactions: textminig, experiments, databases, and co-expression; with high confidence) we found that all coded proteins of the 6 candidates genes interact to each other, except the TAC1 protein (Figure 4A).

**Figure 4.**
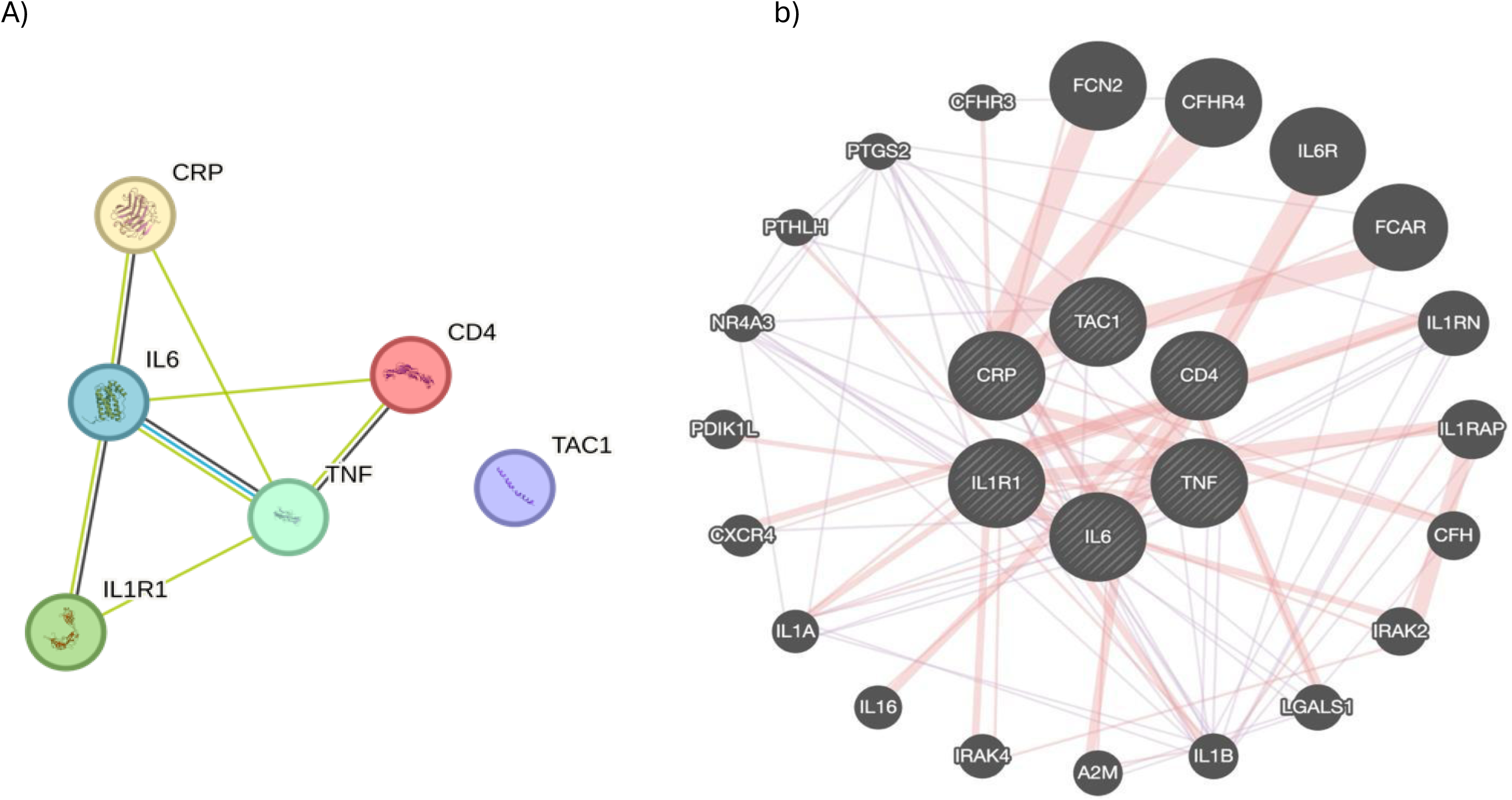
Protein-protein and gene-gene interaction network based on the genes directly modulated by BP drug. A) protein-protein interaction network; B) gene-gene interaction network.

Complementary to the protein-protein interaction analysis, we also investigated the interaction between genes by using the software GENEmania (Warde-Farley, Donaldson et al. 2010). Our analysis indicated that all 6 genes present interactions between them. For example, *TAC1* gene interacts with *IL1R1*, and the genes *IL6* and *TNF* are co-expressed (Figure 4B).

Together, our findings based on tissued expression analysis and network interaction studies indicated that the genes directly modulated by BP are all, at some point, interacting to each other, and thus, the phenotypes associated with each gene can be also affected by the others.

Moreover, EMAP closely resembles the atrophic form of AMD than its neovascular form (Chouraqui, Crincoli et al. 2023). In atrophic AMD, unlike wet AMD, there is greater oxidative stress, mitochondrial dysfunction, and autophagy (Ambati and Fowler 2012, Bowes Rickman, Farsiu et al. 2013). In our analysis, we found that the BP-affected genes *IL6* and *TNF* are related to all of these factors through endoplasmic reticulum stress (Chen, Shi et al. 2023).

Finally, all the genes that are directly modulated by BP are associated with neurodegeneration and auto-immune response, suggesting that long term usage of BP can also drive to the development of EMAP.

Thus, our study provided, for the first time, independent silico molecular data of gene directly modulated by BP that involved in several mechanisms of retinal degeneration, that can lead to the pathophysiological mechanisms of EMAP occurrence.

## Acknowledgement

RHH is supported by Fundação Araucária (grant #FA092016, FA, Paraná, Brazil), National Council for Scientific and Technological Development (grant #402773/2022-5 and grant #311438/2022-9, CNPq, Brazil).

## Conflict of Interests

All authors declare no conflict of interests.

## Supplementary Figure Legends

Figure S1 – Gene expression profile of the CD4 gene in different organs of the human body.

Figure S2 – Gene expression profile of the CRP gene in different organs of the human body.

Figure S3 – Gene expression profile of the IL1R1 gene in different organs of the human body.

Figure S4 – Gene expression profile of the IL6 gene in different organs of the human body.

Figure S5 – Gene expression profile of the TAC1 gene in different organs of the human body.

Figure S6 – Gene expression profile of the TNF gene in different organs of the human body.

## Notes

### Competing Interest Statement

The authors have declared no competing interest.

## References

Ambati, J. and B. J. Fowler (2012). “Mechanisms of age-related macular degeneration.” Neuron 75(1): 26–39.

Antropoli, A., L. Bianco, C. Condroyer, A. Antonio, J. Navarro, D. Dagostinoz, A. Benadji, J.-A. Sahel, C. Zeitz and I. Audo (2024). “Extensive Macular Atrophy with Pseudodrusen-like appearance: Progression Kinetics and Late-Stage Findings.” Ophthalmology 131(10): 1175–1184.

Baek, S. M., S. Y. Yu, Y. Son and H. S. Hong (2016). “Substance P promotes the recovery of oxidative stress-damaged retinal pigmented epithelial cells by modulating Akt/GSK-3β signaling.” Mol Vis 22: 1015–1023.

Blake, G. J. and P. M. Ridker (2002). “Inflammatory bio-markers and cardiovascular risk prediction.” J Intern Med 252(4): 283–294.

Bowes Rickman, C., S. Farsiu, C. A. Toth and M. Klingeborn (2013). “Dry age-related macular degeneration: mechanisms, therapeutic targets, and imaging.” Invest Ophthalmol Vis Sci 54(14): Orsf68-80.

Bray, J. J., S. Thompson, S. Seitler, S. A. Ali, J. Yiu, M. Salehi, M. Ahmad, F. Pelone, H. Gashau, F. Shokraneh, N. Ahmed, M. Cassandra, E. Marijon, D. S. Celermajer and R. Providencia (2024). “Long-term antibiotic prophylaxis for prevention of rheumatic fever recurrence and progression to rheumatic heart disease.” Cochrane Database Syst Rev 9(9): Cd015779.

Carlà, M. M., F. Giannuzzi, F. Boselli, E. Crincoli and S. Rizzo (2024). “Extensive macular atrophy with pseudodrusen-like appearance: comprehensive review of the literature.” Graefes Arch Clin Exp Ophthalmol 262(10): 3085–3097.

Chen, X., C. Shi, M. He, S. Xiong and X. Xia (2023). “Endoplasmic reticulum stress: molecular mechanism and therapeutic targets.” Signal Transduct Target Ther 8(1): 352.

Chirco, K. R. and L. A. Potempa (2018). “C-Reactive Protein As a Mediator of Complement Activation and Inflammatory Signaling in Age-Related Macular Degeneration.” Front Immunol 9: 539.

Chouraqui, M., E. Crincoli, A. Miere, I. A. Meunier and E. H. Souied (2023). “Deep learning model for automatic differentiation of EMAP from AMD in macular atrophy.” Sci Rep 13(1): 20354.

Douillard, A., M. C. Picot, C. Delcourt, A. Lacroux, X. Zanlonghi, B. Puech, S. Defoort-Dhelemmes, I. Drumare, E. Jozefowicz, B. Bocquet, C. Baudoin, N. Al-Dain Marzouka, S. Perez-Roustit, S. Arsène, V. Gissot, F. Devin, C. Arndt, B. Wolff, M. Mauget-Faÿsse, M. Quaranta, T. Mura, D. Deplanque, H. Oubraham, S. Y. Cohen, P. Gastaud, O. Zambrowsky, C. Creuzot-Garcher, S. Mohand Saïd, R. Blanco Garavito, E. Souied, J. A. Sahel, I. Audo, C. Hamel and I. Meunier (2016). “Clinical Characteristics and Risk Factors of Extensive Macular Atrophy with Pseudodrusen: The EMAP Case-Control National Clinical Trial.” Ophthalmology 123(9): 1865–1873.

Falvo, J. V., A. V. Tsytsykova and A. E. Goldfeld (2010). “Transcriptional control of the TNF gene.” Curr Dir Autoimmun 11: 27–60.

Hamel, C. P., I. Meunier, C. Arndt, S. Ben Salah, S. Lopez, C. Bazalgette, C. Bazalgette, X. Zanlonghi, B. Arnaud, S. Defoort-Dellhemmes and B. Puech (2009). “Extensive macular atrophy with pseudodrusen-like appearance: a new clinical entity.” Am J Ophthalmol 147(4): 609–620.

Kilpatrick, J. M. and J. E. Volanakis (1991). “Molecular genetics, structure, and function of C-reactive protein.” Immunol Res 10(1): 43–53.

Kim, S., J. Chen, T. Cheng, A. Gindulyte, J. He, S. He, Q. Li, B. A. Shoemaker, P. A. Thiessen, B. Yu, L. Zaslavsky, J. Zhang and E. E. Bolton (2025). “PubChem 2025 update.” Nucleic Acids Res 53(D1): D1516–d1525.

Klettner, A., A. Brinkmann, K. Winkelmann, T. Käckenmeister, J. Hildebrandt and J. Roider (2020). “Effect of long-term inflammation on viability and function of RPE cells.” Exp Eye Res 200: 108214.

Kutty, R. K., W. Samuel, K. Boyce, A. Cherukuri, T. Duncan, C. Jaworski, C. N. Nagineni and T. M. Redmond (2016). “Proinflammatory cytokines decrease the expression of genes critical for RPE function.” Mol Vis 22: 1156–1168.

Lueck, K., M. Hennig, A. Lommatzsch, D. Pauleikhoff and S. Wasmuth (2012). “Complement and UV-irradiated photoreceptor outer segments increase the cytokine secretion by retinal pigment epithelial cells.” Invest Ophthalmol Vis Sci 53(3): 1406–1413.

Moreira-Neto, C. A., R. A. Schmidt Andujar, J. C. T. Chao, H. Vasconcelos, F. E. E. Alves, G. D. Rodrigues, B. Hirt, J. Arana, E. C. Souza, A. Maia, J. M. F. Sallum and C. A. Moreira, Jr. (2024). “Rheumatic fever and long-term use of benzathine penicillin as possible risk factors for extensive macular atrophy with pseudodrusen in a Brazilian cohort.” Int J Retina Vitreous 10(1): 75.

Mosley, B., D. L. Urdal, K. S. Prickett, A. Larsen, D. Cosman, P. J. Conlon, S. Gillis and S. K. Dower (1987). “The interleukin-1 receptor binds the human interleukin-1 alpha precursor but not the interleukin-1 beta precursor.” J Biol Chem 262(7): 2941–2944.

Nielsen, M. K., Y. Subhi, M. Falk, A. Singh, T. L. Sørensen, M. H. Nissen and C. Faber (2023). “Complement factor H Y402H polymorphism results in diminishing CD4(+) T cells and increasing C-reactive protein in plasma.” Sci Rep 13(1): 19414.

Peng, L., G. O. Agogo, J. Guo and M. Yan (2019). “Substance P and fibrotic diseases.” Neuropeptides 76: 101941.

Schmidt-Arras, D. and S. Rose-John (2016). “IL-6 pathway in the liver: From physiopathology to therapy.” J Hepatol 64(6): 1403–1415.

Sjöstedt, E., W. Zhong, L. Fagerberg, M. Karlsson, N. Mitsios, C. Adori, P. Oksvold, F. Edfors, A. Limiszewska, F. Hikmet, J. Huang, Y. Du, L. Lin, Z. Dong, L. Yang, X. Liu, H. Jiang, X. Xu, J. Wang, H. Yang, L. Bolund, A. Mardinoglu, C. Zhang, K. von Feilitzen, C. Lindskog, F. Pontén, Y. Luo, T. Hökfelt, M. Uhlén and J. Mulder (2020). “An atlas of the protein-coding genes in the human, pig, and mouse brain.” Science 367(6482).

Subhi, Y., M. Krogh Nielsen, C. R. Molbech, A. Oishi, A. Singh, M. H. Nissen and T. L. Sørensen (2019). “Plasma markers of chronic low-grade inflammation in polypoidal choroidal vasculopathy and neovascular age-related macular degeneration.” Acta Ophthalmol 97(1): 99–106.

Szklarczyk, D., R. Kirsch, M. Koutrouli, K. Nastou, F. Mehryary, R. Hachilif, A. L. Gable, T. Fang, N. T. Doncheva, S. Pyysalo, P. Bork, L. J. Jensen and C. von Mering (2023). “The STRING database in 2023: protein-protein association networks and functional enrichment analyses for any sequenced genome of interest.” Nucleic Acids Res 51(D1): D638–d646.

Todd, L., I. Palazzo, L. Suarez, X. Liu, L. Volkov, T. V. Hoang, W. A. Campbell, S. Blackshaw, N. Quan and A. J. Fischer (2019). “Reactive microglia and IL1β/IL-1R1-signaling mediate neuroprotection in excitotoxin-damaged mouse retina.” J Neuroinflammation 16(1): 118.

Touhami, S., F. Beguier, S. Augustin, H. Charles-Messance, L. Vignaud, E. F. Nandrot, S. Reichman, V. Forster, T. Mathis, J. A. Sahel, B. Bodaghi, X. Guillonneau and F. Sennlaub (2018). “Chronic exposure to tumor necrosis factor alpha induces retinal pigment epithelium cell dedifferentiation.” J Neuroinflammation 15(1): 85.

Warde-Farley, D., S. L. Donaldson, O. Comes, K. Zuberi, R. Badrawi, P. Chao, M. Franz, C. Grouios, F. Kazi, C. T. Lopes, A. Maitland, S. Mostafavi, J. Montojo, Q. Shao, G. Wright, G. D. Bader and Q. Morris (2010). “The GeneMANIA prediction server: biological network integration for gene prioritization and predicting gene function.” Nucleic Acids Res 38(Web Server issue): W214-220.

Wolf, J., S. Boneva, A. Schlecht, T. Lapp, C. Auw-Haedrich, W. Lagrèze, H. Agostini, T. Reinhard, G. Schlunck and C. Lange (2022). “The Human Eye Transcriptome Atlas: A searchable comparative transcriptome database for healthy and diseased human eye tissue.” Genomics 114(2): 110286.

Xiao, R., C. Lei, Y. Zhang and M. Zhang (2023). “Interleukin-6 in retinal diseases: From pathogenesis to therapy.” Exp Eye Res 233: 109556.

Zeitlmann, L., P. Sirim, E. Kremmer and W. Kolanus (2001). “Cloning of ACP33 as a novel intracellular ligand of CD4.” J Biol Chem 276(12): 9123–9132.

Zeng, Y., X. Yin, C. Chen and Y. Xing (2021). “Identification of Diagnostic Biomarkers and Their Correlation with Immune Infiltration in Age-Related Macular Degeneration.” Diagnostics (Basel) 11(6).

