## Supplementary material for "Rapid Report: Retinal gene and pathway modulation by benzathine penicillin: A potential link to Extensive Macular Atrophy with Pseudodrusen (EMAP) and neurodegeneration": Sup. Figures

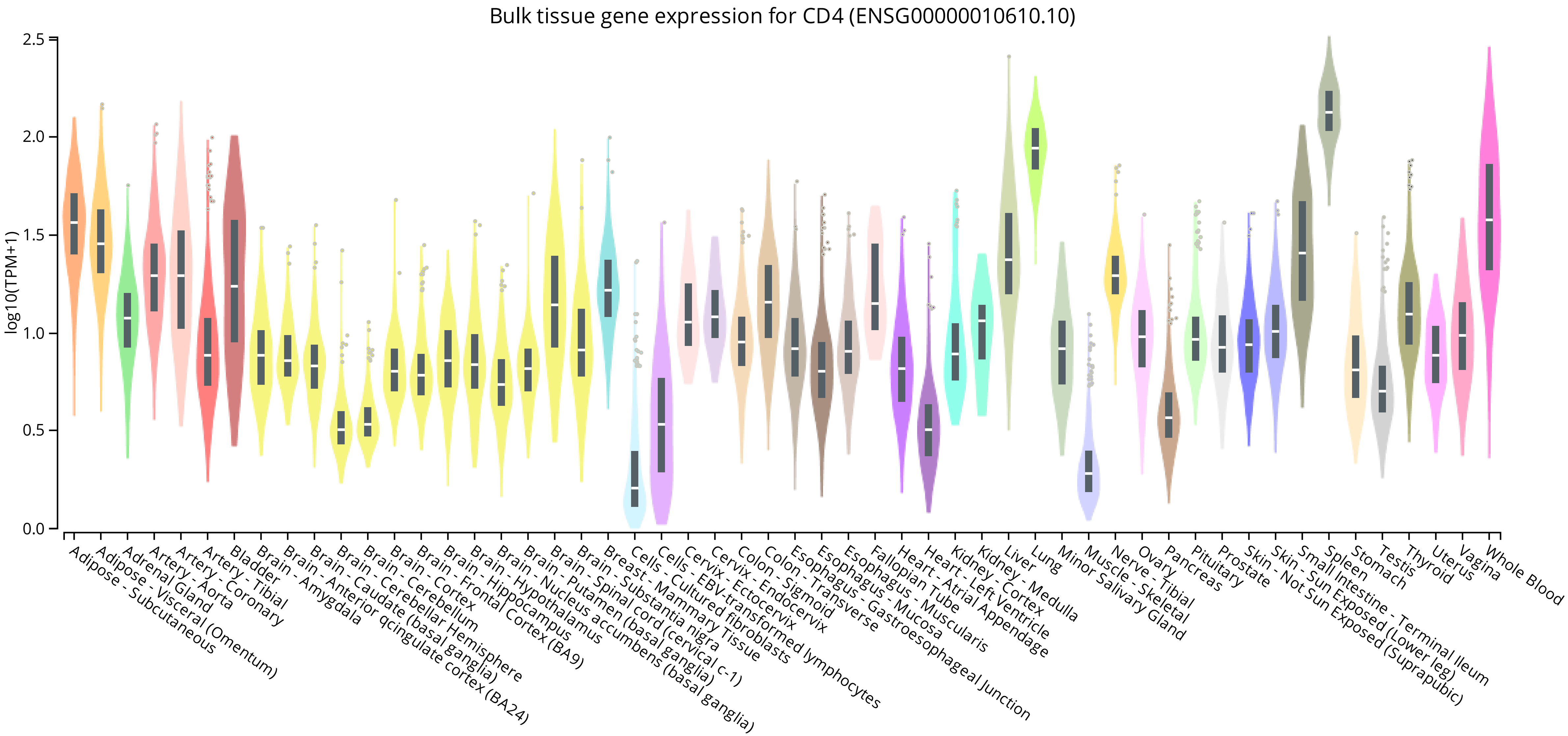


Figure S1

Shinsato & Herai, 2025

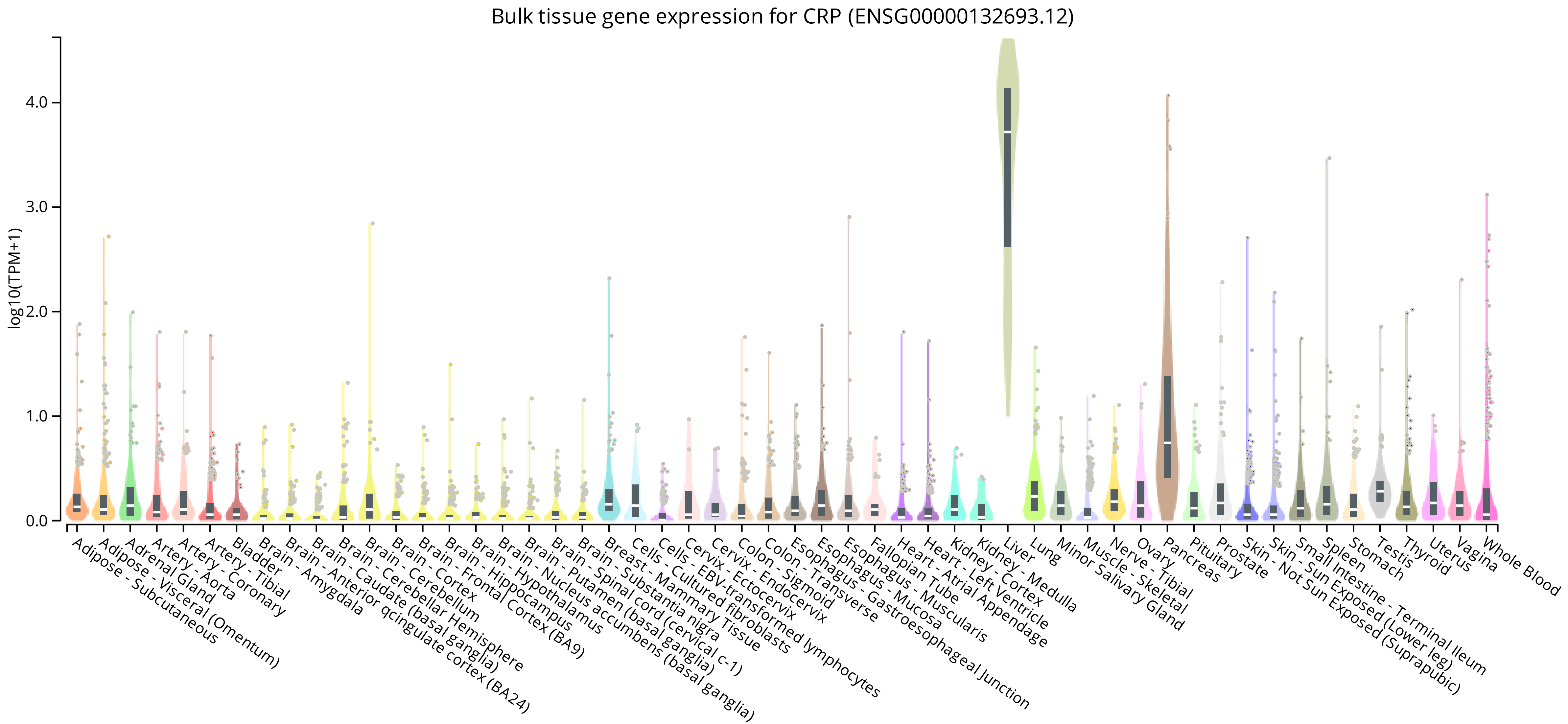


Figure S2

Shinsato & Herai, 2025

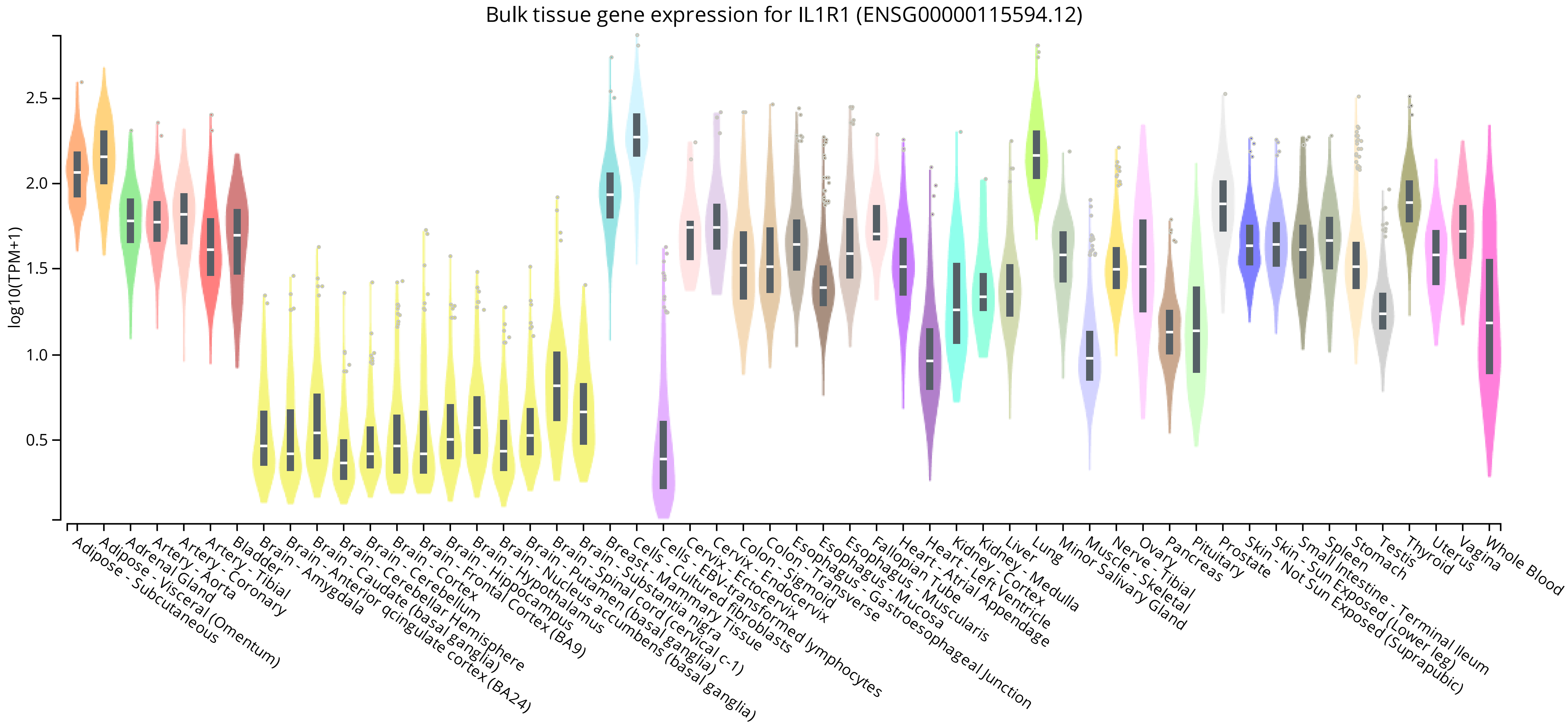


Figure S3

Shinsato & Herai, 2025

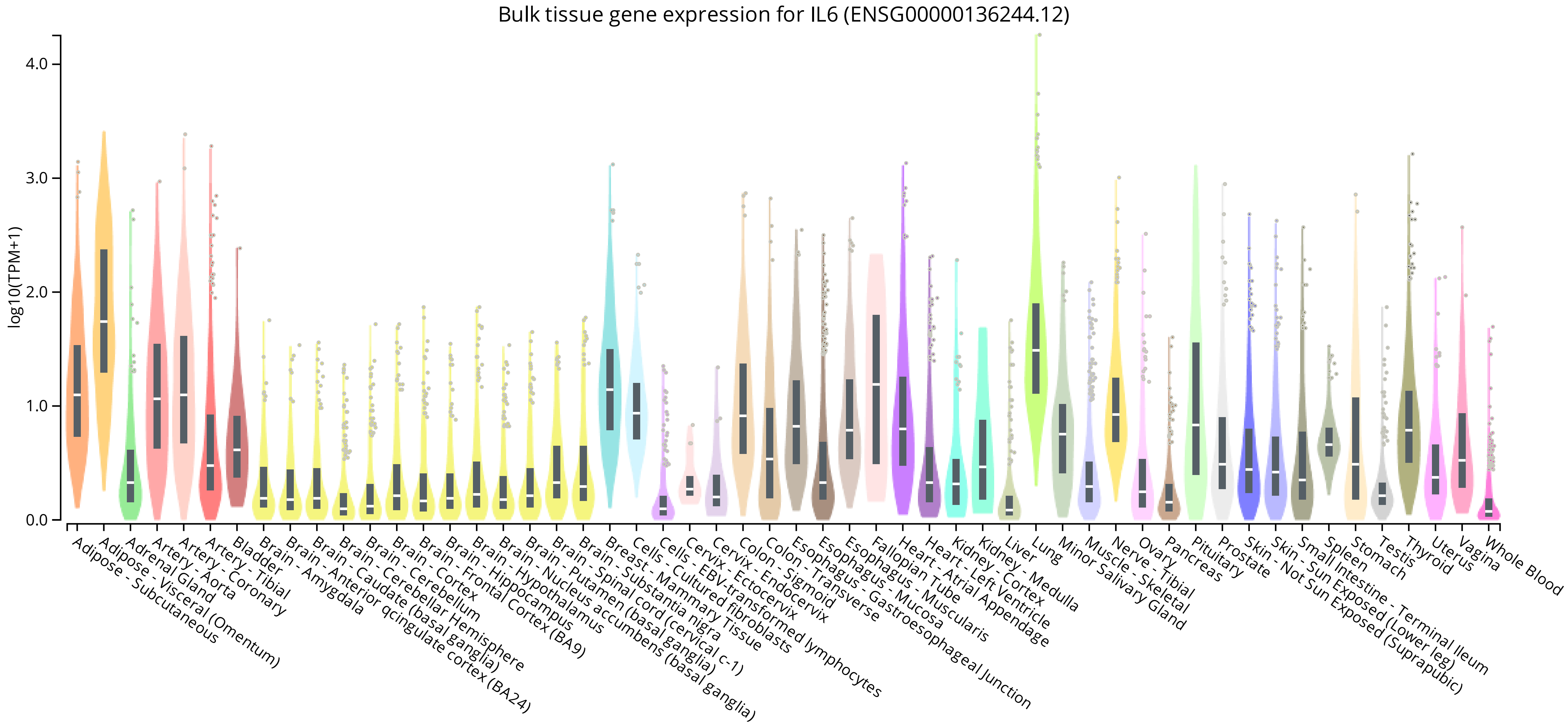


Figure S4

Shinsato & Herai, 2025

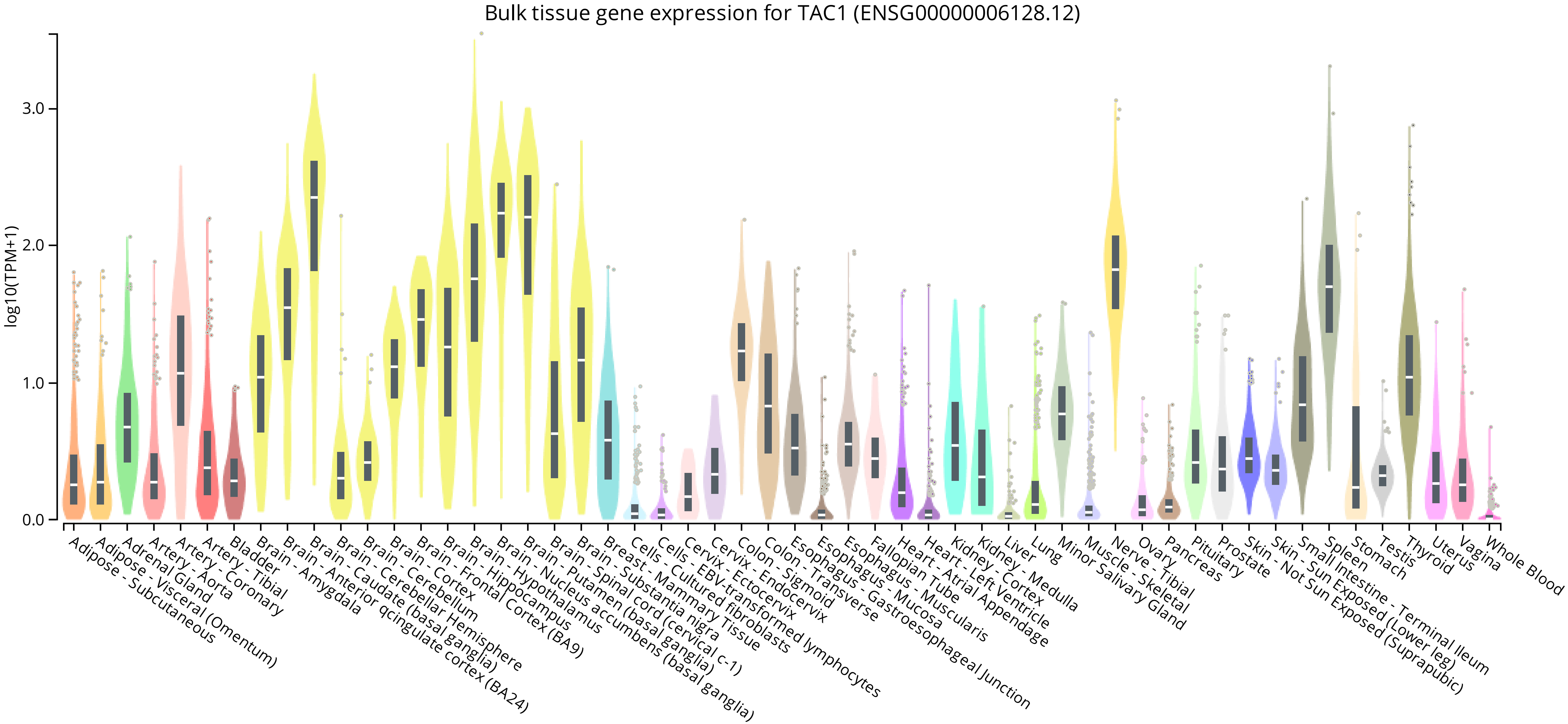


Figure S5

Shinsato & Herai, 2025

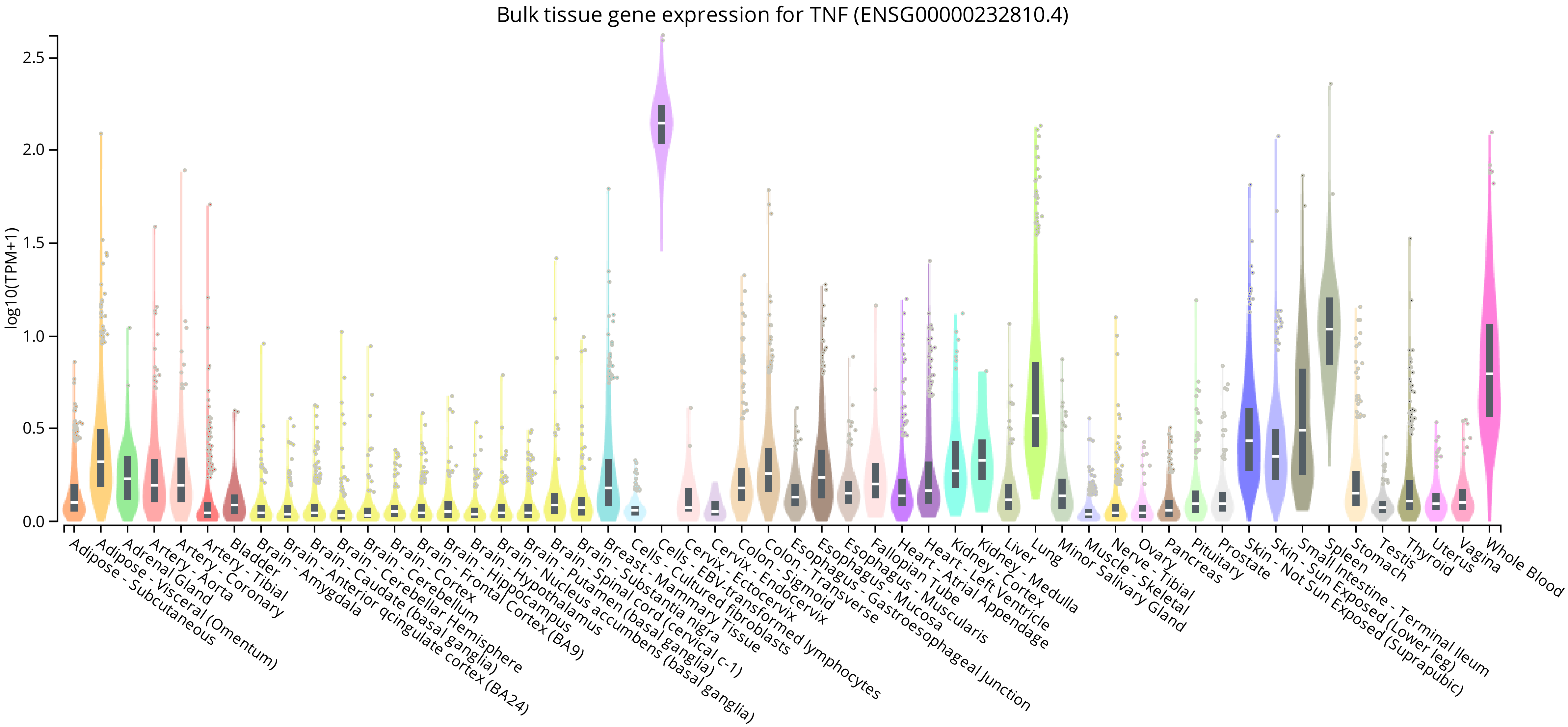


Figure S6

Shinsato & Herai, 2025
